# Comparison of transcriptional responses and metabolic alterations in three multidrug resistant model microorganisms, *Staphylococcus aureus* ATCC BAA-39, *Escherichia coli* ATCC BAA-196 and *Acinetobacter baumannii* ATCC BAA-1790, on exposure to iodine-containing nano-micelle drug FS-1

**DOI:** 10.1101/2020.09.01.278945

**Authors:** Ilya S. Korotetskiy, Sergey V. Shilov, Tatyana V. Kuznetsova, Aleksandr I. Ilin, Monique Joubert, Setshaba Taukobong, Oleg N. Reva

## Abstract

Iodine is one of the oldest antimicrobial agents. Till now there have been no reports on acquiring resistance to iodine. Recent studies showed promising results on application of iodine-containing nano-micelles, FS-1, against antibiotic resistant pathogens as a supplement to antibiotic therapy. The mechanisms of the action, however, remain unclear. The aim of this study was to perform a holistic analysis and comparison of gene regulation in three phylogenetically distant multidrug resistant reference strains representing pathogens associated with nosocomial infections from the ATCC culture collection: *Escherichia coli* BAA-196, *Staphylococcus aureus* BAA-39 and *Acinetobacter baumannii* BAA-1790. These cultures were treated by a 5 min exposure to sublethal concentrations of the iodine-containing drug FS-1 applied in the late lagging and the mid of the logarithmic growth phases. Complete genome sequences of these strains were obtained in the previous studies. Gene regulation was studied by total RNA extraction and Ion Torrent sequencing followed by mapping the RNA reads against the reference genome sequences and statistical processing of read counts using the DESeq2 algorithm. It was found that the treatment of bacteria with FS-1 profoundly affected the expression of many genes involved in central metabolic pathways; however, alterations of the gene expression profiles were species-specific and depended on the growth phase. Disruption of respiratory electron-transfer membrane complexes, increased penetrability of bacterial cell walls, osmotic and oxidative stresses leading to DNA damaging were the major factors influencing the treated bacteria.

**IMPORTANCE:** Infections caused by antibiotic resistant bacteria threaten the public health worldwide. Combinatorial therapy when antibiotics are administrated together with supplementary drugs improving susceptibility of pathogens to the regular antibiotics is considered as a promising way to overcome this problem. An induction of antibiotic resistance reversion by the iodine-containing nano-micelle drug FS-1 has been reported recently. This drug is currently under clinical trials in Kazakhstan against multidrug resistant tuberculosis. The effects of released iodine on metabolic and regulatory processes in bacterial cells remain unexplored. The current work provides an insight into gene regulation in the antibiotic resistant nosocomial reference strains treated with iodine-containing nanoparticles. This study sheds light on unexplored bioactivities of iodine and the mechanisms of its antibacterial effect when applied in sublethal concentrations. This knowledge will aid in the future design of new drugs against antibiotic resistant infections.

## Introduction

Iodine was discovered by French chemist Bernard Courtois in 1811. Since then it has been used as one of the most successful disinfectants, which antimicrobial activity has never been compromised by any acquired or natural resistance. The history of discovery and a comprehensive overview of the use of iodine in medicine was published in 1961 on the 150-year anniversary of the discovery of this chemical element [1]. Iodine has, however, significantly lost its ground in medicine after the discovery of antibiotics by Alexander Fleming in 1928 [2, 3]. Although originally developed for treating human infectious diseases, remarkable antimicrobial properties of antibiotics have spread them to a broad application in farming, agroindustry and aquaculture [4]. The misuse and overuse of antibiotics has led to a strong selective pressure towards emerging and wide distribution of antibiotic resistant pathogens threatening the public health systems worldwide [2, 4]. Eventually, the situation with the antibiotic resistance and the problems with development of new antibiotics for an affordable price came to a global crisis [5–8]. An urgent need in alternative approaches to cope with the problem of antibiotic resistance is generally recognized [9, 10]. It is of great importance to devise new strategies that focus on limiting, redirecting and/or reversing the antibiotic resistance in bacterial.

Surprisingly, despite having used iodine in medicine for over 200 years, the effect of iodine and its derivative compounds on metabolic processes and gene expression in living organisms remains basically unexplored except for numerous studies on iodine-rich thyroid hormones. A reason for this may be that iodine was used in medicine mostly for topical treatment of wounds or in composition of industrial disinfectants designed for a rapid eradication of germs leaving no time for the treated bacteria to respond to the treatment either by gene regulation or metabolic adaptation. Only recently, several published works demonstrated rather complex mechanisms of the action of iodine on eukaryotic cells and gene expression patterns [11–13]. An interesting fact was published by Klebanoff in 1967 [14] on the release of iodinated organic substances by leukocytes at spotted infections that suggested a possible natural therapeutic effect of iodine in concentrations much lower than that in the disinfectants. This discovery had no continuation in any further studies probably because the iodine-based antimicrobials looked in the flourishing days of the antibiotic era as an unpromising archaic aftermath of the 19^th^ century medicine. However, the post-antibiotic era calls us to reconsider applicability of previously abandoned anti-infection remedies [15].

Iodine-containing drugs seems to provide a viable option to overcome the antibiotic resistance problem, since they have strong antimicrobial properties and no acquired resistance to iodine has been reported [16–18]. Molecular iodine is an effective mild and nontoxic Lewis acid catalyst [19] of great potential for synthesis of various macromolecular complexes [20] that can easily penetrate cells of microorganisms [21] and pass through bi-lipid layers of cell membranes [11]. Since iodine can easily penetrate cell membranes, its application may be efficient against intracellular infections. This concept was implemented in a new drug, FS-1, developed as a new medicine against multidrug resistant tuberculosis [21, 22]. FS-1 contains iodine molecules and ions bound to dextrin/oligopeptide nano-micelles. General composition of FS-1 particles is shown below:

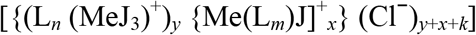

where L – dextrin-polypeptide ligand; Me – Li/Mg ions; *n*, *m*, *x*, *y* and *k* – variable integers ≥ 1. Molecular mass of the micelles is in the range of 30–300 kD.

FS-1 has passed both the preclinical and clinical trials (www.clinicaltrials.gov, accession number NCT02607449) and was approved in Kazakhstan as a supplementary drug to be administrated with the common antibiotics used against multidrug resistant (MDR) and extensively drug resistant (XDR) tuberculosis. It was demonstrated that the application of FS-1 increased the susceptibility of the treated bacteria to antibiotics [22]; however, molecular mechanisms of this action remain unclear. As the use of multidrug resistant *M. tuberculosis* isolates for further studies was technically challenged, the study of the action of iodine-containing nano-micelles on antibiotic resistant bacteria was continued using ATCC collection strains *Escherichia coli* BAA-196, *Staphylococcus aureus* BAA-39 and *Acinetobacter baumannii* BAA-1790. Complete genome sequences of these microorganisms were obtained using SMRT PacBio sequencing for two variants of the strains: the initial culture and the culture grown with sub-bactericidal concentrations of FS-1 in 10 daily passages [23–25]. Global genetic variations, epigenetic modifications and altered gene regulation in *S. aureus* BAA-39 and *E. coli* BAA-196 cultivated with FS-1 were analysed in recent publications [26–28]. The aim of the current study was to perform a systemic analysis of gene expression and metabolic alteration in these three selected multidrug resistant pathogens affected by a 5 min exposure to the sublethal dose of the iodine-containing drug FS-1 in the late lagging and the mid of the logarithmic growth phases. Our expectation is that this study will shed light on unexplored bioactivities of iodine and that the comprehensive knowledge of metabolic and regulatory pathways affected by iodine-containing nano-micelles will aid in future design of new drugs against multidrug resistant infections.

## Results

The effect of a 5 min exposure to FS-1 on gene expression was analysed on three model microorganisms: *E. coli* ATCC BAA-196, *S. aureus* ATCC BAA-39 and *A. baumannii* ATCC BAA-1790. Total RNA extracts were converted to cDNA and sequenced by the Ion Torrent technology. Then the DNA reads were mapped against the complete genome sequences of these microorganisms. The drug FS-1 was applied at the end of the lagging growth phase (Lag-experiment) and in the mid of the logarithmic growth phase (Log-experiment). Growth curves estimated for the model microorganisms are shown in Supplementary Fig. S1. Each experiment was repeated three times (with *E. coli*) or six times (with *S. aureus* and *A. baumannii*) to accumulate enough reads and repeats for achieving a statistical confidence for the differentially expressed genes (Table 1). The counts of reads overlapping individual genes were normalized by the program Bioconducter DESeq2 and the statistical parameters of differential gene expression were calculated.

**Table 1.**
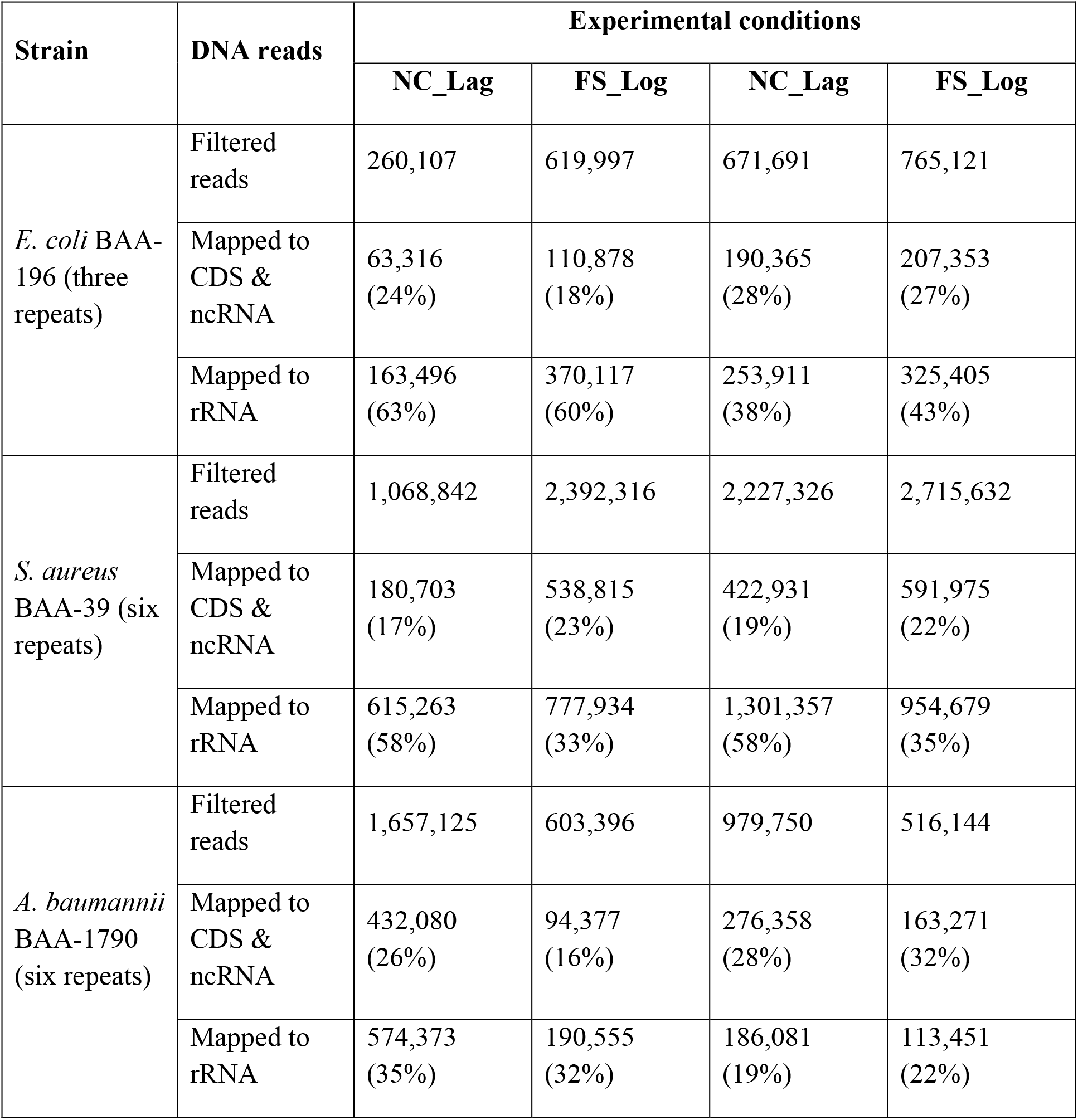
Numbers of DNA reads after quality filtering and numbers of reads mapped against coding genes (CDS & ncRNA) and rRNA genes. Negative control (NC) and experimental (FS) samples obtained from the Lag and Log experiments are indicated in the column titles.

| Strain | DNA reads | Experimental conditions |  |  |  |
| --- | --- | --- | --- | --- | --- |
|  |  | NC_Lag | FS_Log | NC_Lag | FS_Log |
| *E. coli* BAA-196 (three repeats) | Filtered reads | 260,107 | 619,997 | 671,691 | 765,121 |
|  | Mapped to CDS & ncRNA | 63,316 (24%) | 110,878 (18%) | 190,365 (28%) | 207,353 (27%) |
|  | Mapped to rRNA | 163,496 (63%) | 370,117 (60%) | 253,911 (38%) | 325,405 (43%) |
| *S. aureus* BAA-39 (six repeats) | Filtered reads | 1,068,842 | 2,392,316 | 2,227,326 | 2,715,632 |
|  | Mapped to CDS & ncRNA | 180,703 (17%) | 538,815 (23%) | 422,931 (19%) | 591,975 (22%) |
|  | Mapped to rRNA | 615,263 (58%) | 777,934 (33%) | 1,301,357 (58%) | 954,679 (35%) |
| *A. baumannii* BAA-1790 (six repeats) | Filtered reads | 1,657,125 | 603,396 | 979,750 | 516,144 |
|  | Mapped to CDS & ncRNA | 432,080 (26%) | 94,377 (16%) | 276,358 (28%) | 163,271 (32%) |
|  | Mapped to rRNA | 574,373 (35%) | 190,555 (32%) | 186,081 (19%) | 113,451 (22%) |

Gene expression differences equal or higher than 2-folds with estimated *p*-values equal or lower than 0.05 were considered as significant and statistically reliable. Numbers of up- and downregulated genes in the Lag and Log growth phases in three model microorganisms are shown in Table 2. Involvement of the differentially expressed genes in metabolic pathways was predicted by the program Pathway Tools v14.0. A summary of regulated metabolic pathways in the three model organisms is shown in Table 3. Also, all differentially expressed genes identified in *E. coli*, *S. aureus* and *A. baumannii* are listed in supplementary Tables S1, S2 and S3, respectively, including involvements of these genes in metabolic pathways predicted by the program Pathway Tools.

**Table 2.** Numbers of regulated genes identified in the three model microorganisms treated with FS-1 in the Lag and Log growth phases.

| Strain | Lag growth phase |  | Log growth phase |  |
| --- | --- | --- | --- | --- |
|  | Up-regulated | Down-regulated | Up-regulated | Down-regulated |
| *E. coli* ATCC BAA-196 | 42 | 78 | 15 | 43 |
| *S. aureus* ATCC BAA-39 | 223 | 230 | 76 | 29 |
| *A. baumannii* ATCC BAA-1790 | 102 | 79 | 78 | 127 |

**Table 3.**
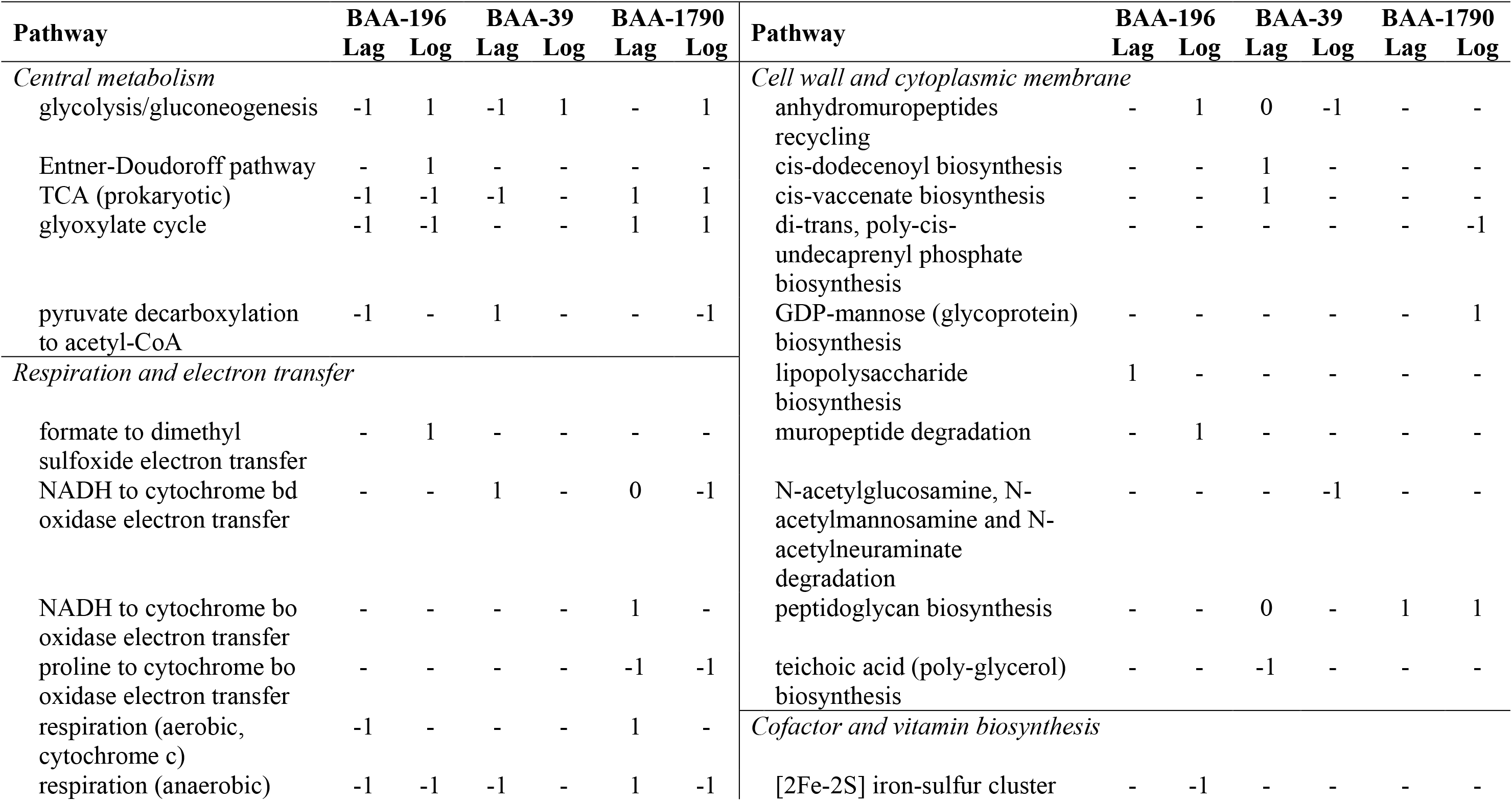
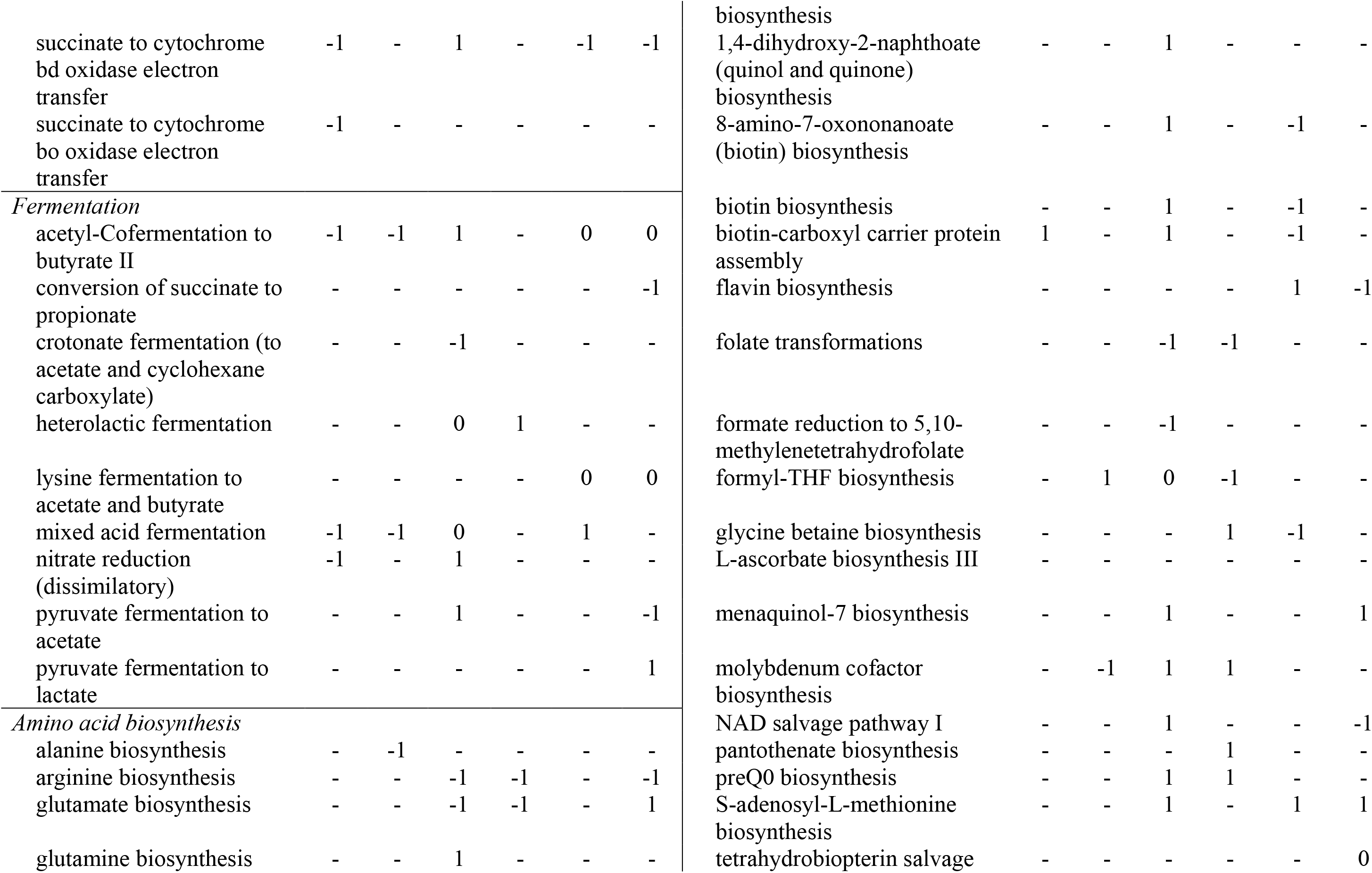
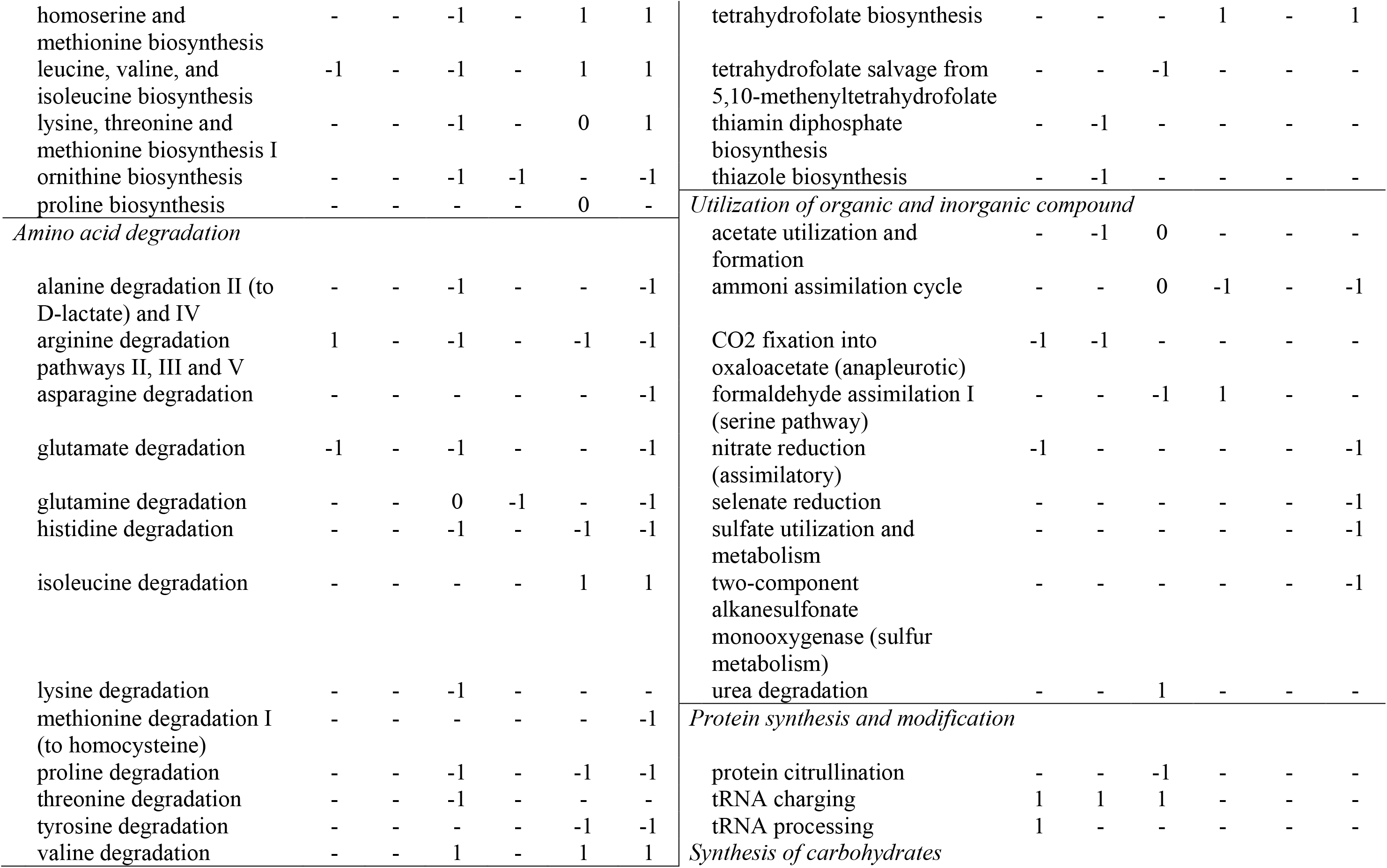
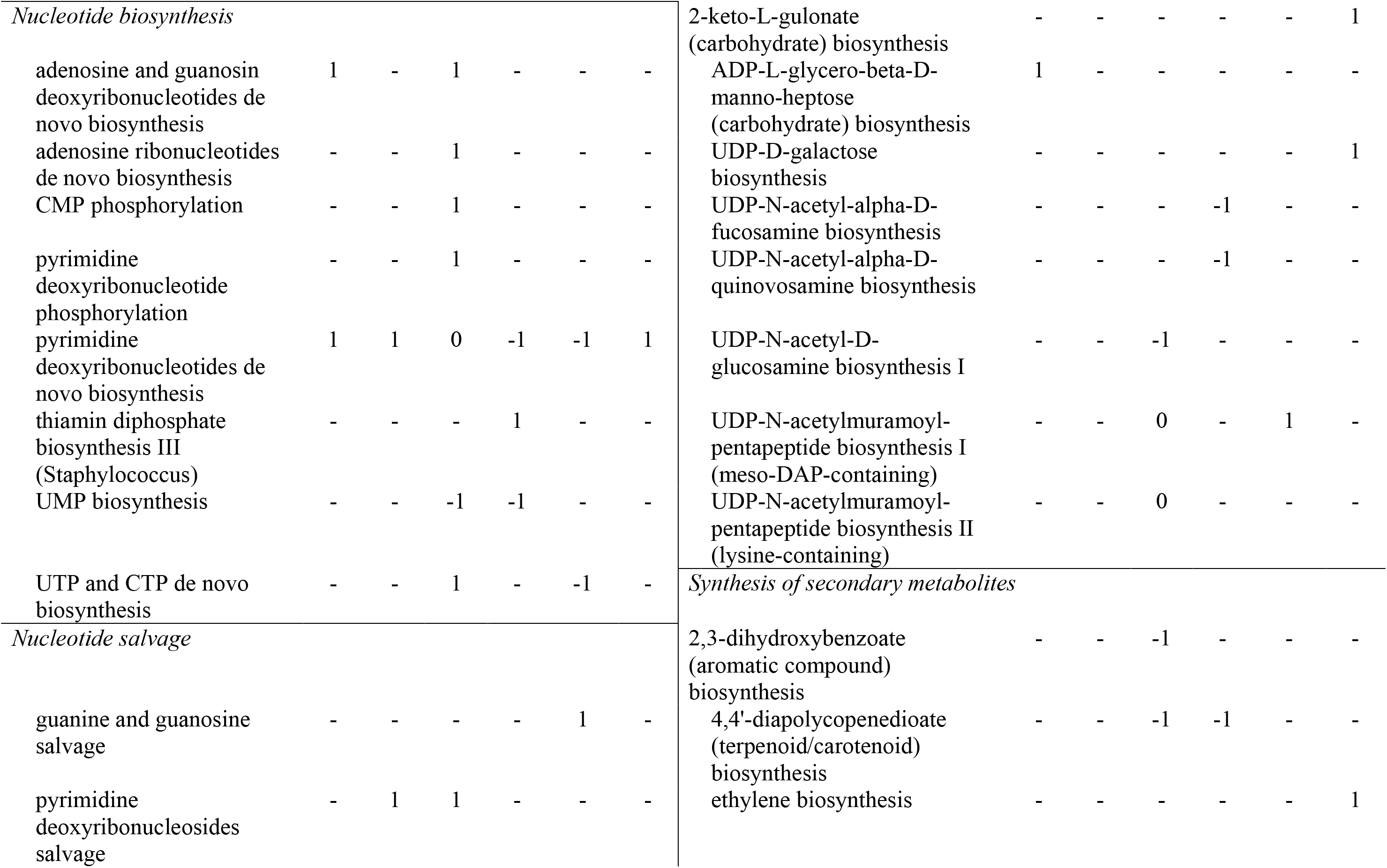
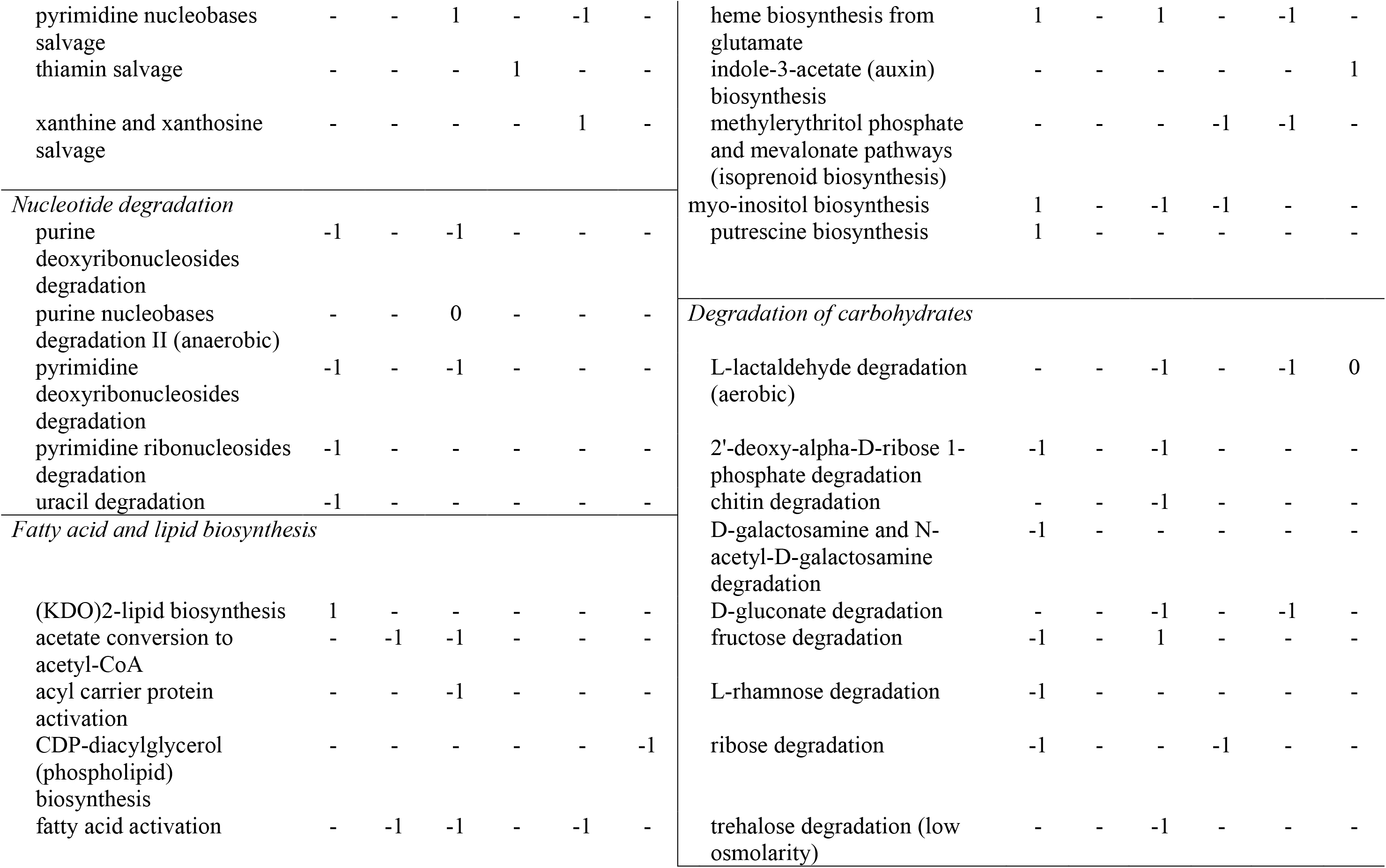
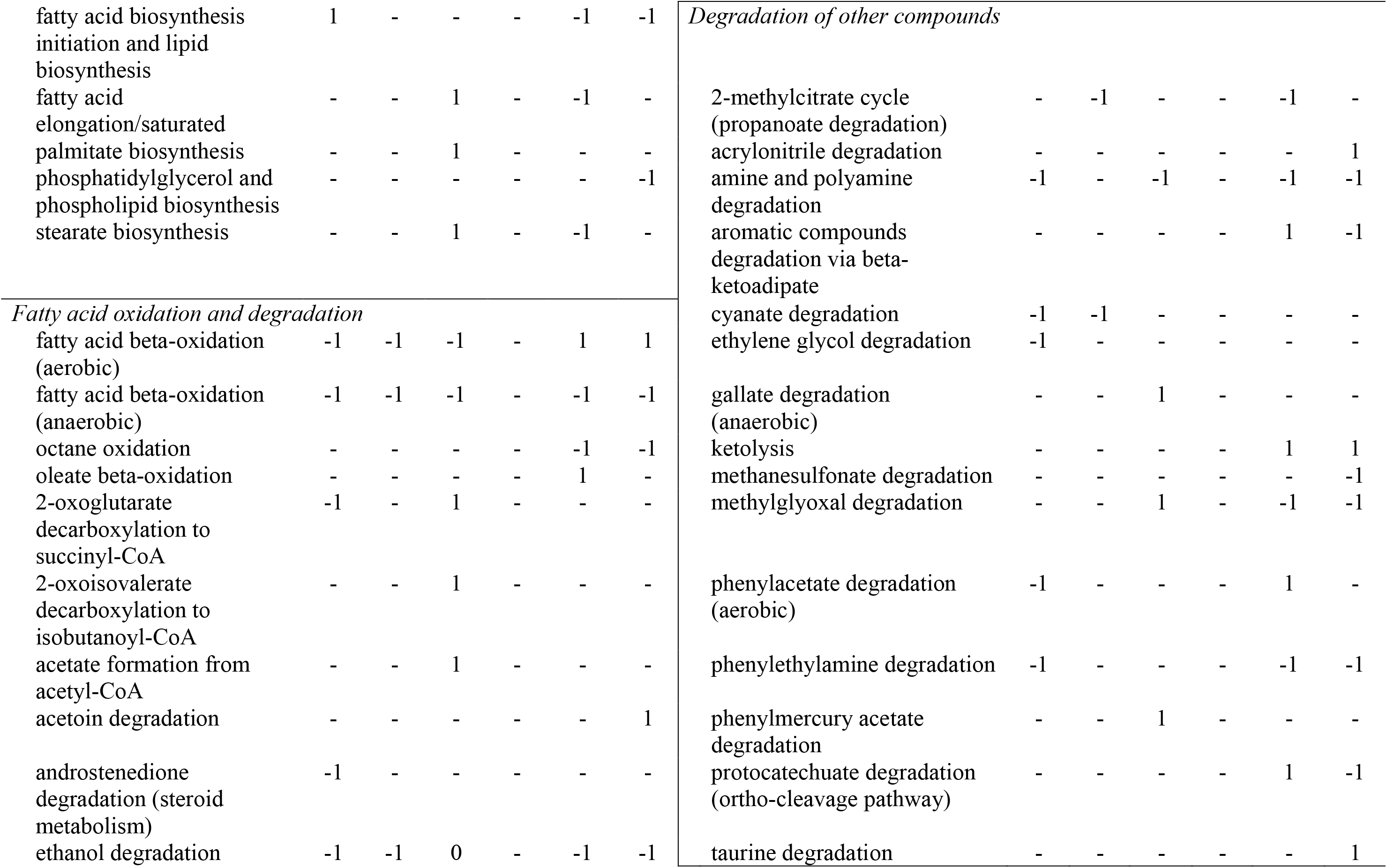
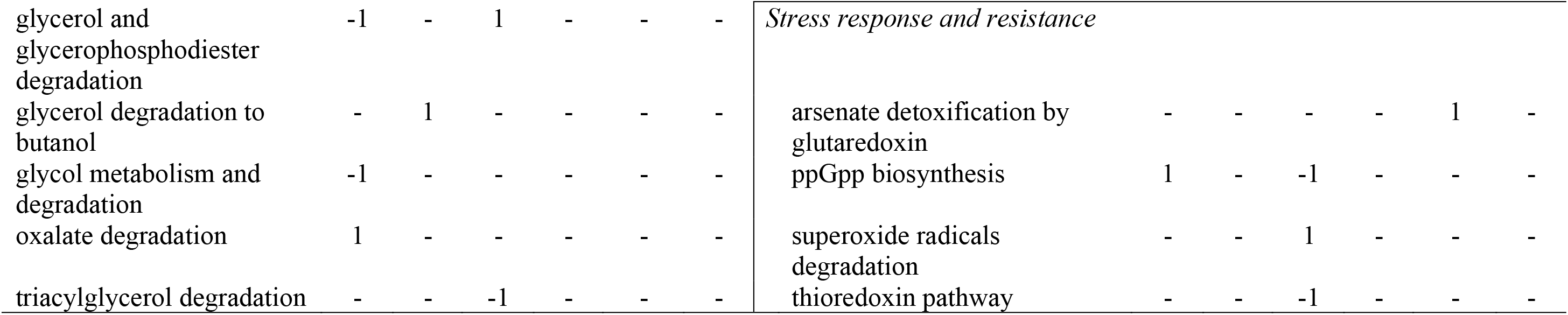
Regulation of metabolic pathways by the treatment with FS-1 of *E. coli* ATCC BAA-196, *S. aureus* ATCC BAA-39 and *A. baumannii* ATCC BAA-1790 at the Lag and Log growth phases as predicted by the program Pathway Tools v14.0 for the regulated genes (see Supplementary Tables S1-S3). Upregulation is depicted by 1; downregulation is depicted by −1; alternative regulation of genes involved in the pathway is depicted by 0.

The gene regulation caused by the treatment with FS-1 at different growth phases showed some level of co-regulation with aggregated Pearson correlation coefficients 0.504, 0.441 and 0.259 calculated at the Lag and Log experimental conditions for all differentially expressed genes identified respectively for *E. coli*, *S. aureus* and *A. baumannii* (Fig. 1). In the case with *A. baumannii*, the correlation was smaller due to an opposite regulation of several genes involved in taurine, sulphate and nitrate transportation and (*tauAB*, *cysA*, *cysP*, *cysW*, *ybaN*), which were significantly downregulated by the treatment with FS-1 in the Lag growth phase but upregulated in the Log phase at the same condition. In *S. aureus*, an alternative regulation was observed for the gene *pyrC* involved in pyrimidine *de novo* biosynthesis.

**Fig. 1.**
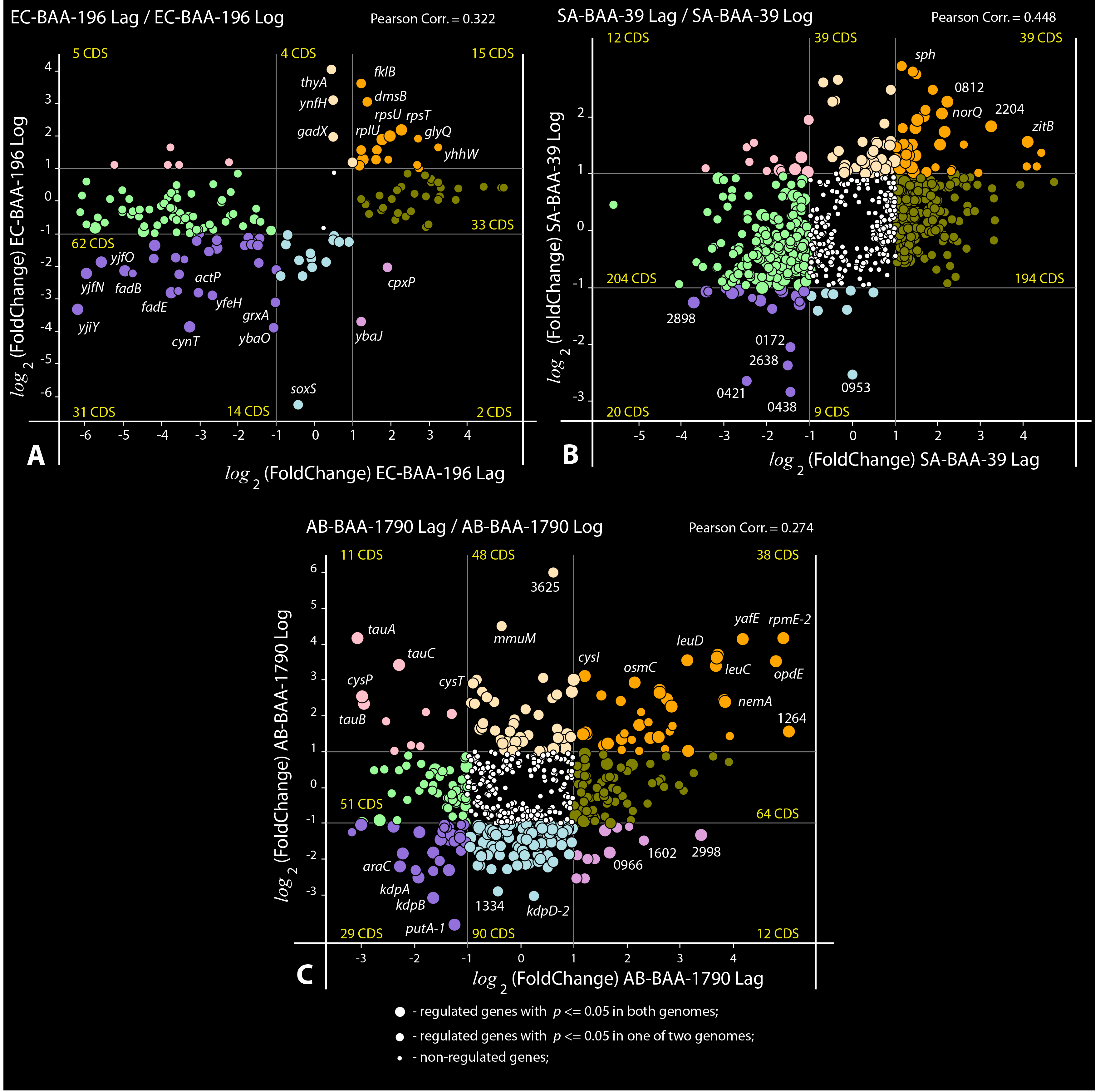
Plots of co-regulation of gene in the Lag and Log growth phases. A) *E. coli* BAA-196 (EC-BAA-196 Lag/Log); B) *S. aureus* BAA-39 (SA-BAA-39 Lag/Log); and C) *A. baumannii* BAA-1790 (AB-BAA-1790 Lag/Log). Circles represent protein coding genes (CDS) plotted according to their negative and positive *log*_2_(fold-change) values calculated in the Lag-experiment (axis X) and Log-experiment (axis Y). The outermost regulated genes are labelled by their names or locus tag numbers. Thin vertical and horizontal lines within the plots separate genes with 2-fold or higher regulation and split the plots into sectors of genes of different categories depending on their coregulation. Numbers of CDS falling to different sectors are shown. Up- and down-coregulated genes, oppositely regulated genes and the genes regulated only in one experiment are depicted by different colours. Statistical reliability of the fold change predictions is depicted by sizes of circles as explained in the legend on the bottom part of the figure. Estimated Pearson correlation coefficients are given on the top of each plot.

### Differential gene expression in *E. coli* ATCC BAA-196

Differential gene expression in *E. coli* treated for 5 min with FS-1 in the Lag and Log growth phases is shown in Supplementary Table S1. The scheme of regulated metabolic pathways is shown in Fig. 2A.

**Fig. 2.**
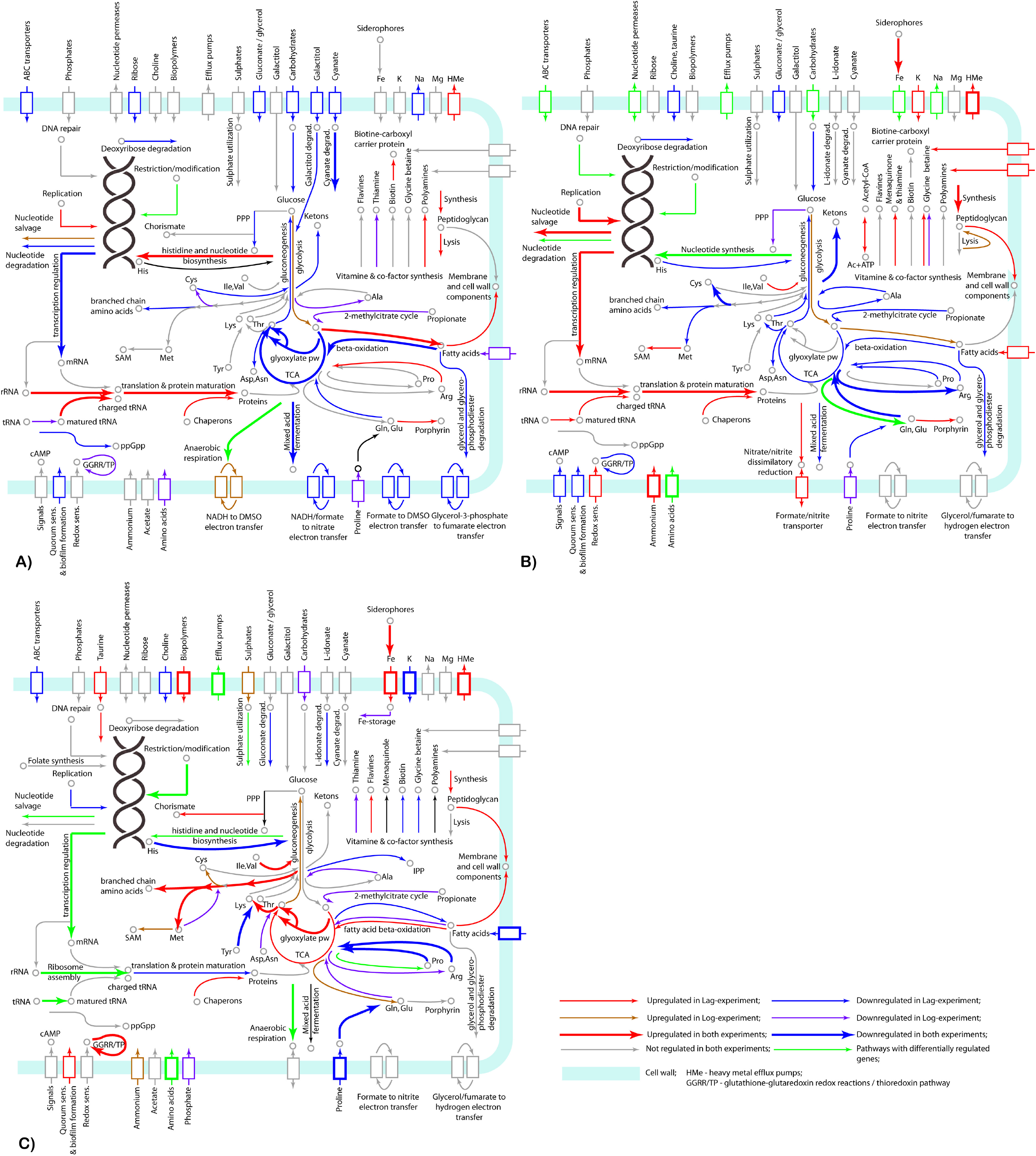
Metabolic pathways affected by the treatment of the model microorganisms A) *E. coli* BAA-196, B) *S. aureus* BAA-39 and C)*A. baumannii* BAA-1790; by the treatment with FS-1 as predicted by the gene regulation analysis in the Lag and Log experiments. Up- and down-regulation of metabolic pathways discovered in both, or only in one experiment, are depicted by arrows of different colours and width as explained in the legend on the right-bottom part of the figure. Cell membrane and cell wall associated proteins are shown by blocks depicted by the same colour scheme. Individual pathways and key compounds are labelled.

The treatment with FS-1 has affected the central metabolism of the bacterium. All genes of the tricarboxylic acid cycle (TCA), *fumA*, *sdhA*, *sdhB*, *sucA*, *sucB* and *sucC*, were strongly downregulated at least in one growth phase. The glyoxylate shunt genes *acnB* and *gltA* were downregulated in the Log phase. Glycolytic gene *pfkA* was 3-fold upregulated in the Lag and Log experiments and another glycolytic gene *tpiA* was 3-fold upregulated in the Log-phase. All these observations indicate a general trend on an activation of glycolysis in the FS-1 treated *E. coli*. Strong suppression of the TCA cycle with the upregulation of glycolysis are indicative for the anaerobiosis [29]. A complete switch to anaerobiosis was reported for this strain cultivated for 10 passages with FS-1 as reported in the previous study [27, 28]. In this study, a strong upregulation of NADH:quinone oxidoreductase *ndh* was observed in both growth phases. This enzyme is important for the aerobic and anaerobic respiration. It participates in transportation of protons from NADH to oxygen through a cascade of H^+^-transporting ubiquinol oxidases CyoABCD (these genes were insignificantly downregulated by FS-1) or CydABXH (insignificantly upregulated), or to nitrates through ubiquinol:nitrate oxidoredutases NarGHI (*narG* was 6.6-fold downregulated, *p* = 0.09). Mixed acid fermentation enzymes (*acnB*, *adhE*, *gltA*, *fdhF*, *frdA*, *frdB*) and all other electron transfer systems were mostly downregulated, namely: formate-to-dimethyl sulfoxide (DMSO) / nitrite / trimethylamine N-oxide (*fdoGH*); glycerol-3-phosphate (G3P)-to-cytochrome *bo* oxidas (*glpD*); G3P-to-fumarate/hydrogen peroxide reductases (*glpAB*, *frdA*); NADH/hydrogen-to-fumarate reductase (*frdA*) and succinate-to-cytochrome *bd* oxidase (*sdhAB*). An only exception was the upregulated NADH-to-DMSO electron transfer system catalysed by a complex of dimethyl sulfoxide reductase DmsABC with menaquinol. In the Log-phase, *dmsB* was 8-fold upregulated. It suggests a possible transition of the FS-1 treated *E. coli* towards anaerobic respiration using DMSO as a terminal electron acceptor.

Many catabolic processes and transmembrane transportation systems were inhibited in the FS-1 treated cells. Downregulation was observed for the genes involved in carbohydrate transportation (*lamB*, *malE*, *malK*, *mglABC*, *treB*) and degradation (*fruK*); acetate/glycolate permease *actP*, galactitol transportation and degradation (*gatYZ*); cyanate transportation and degradation (*cynS*, *cynT*, *cynX*); glycerol and glycero-phosphodiester degradation (*glpABDKO*, *dhaKL*); propanoate degradation through 2-methylcitrate cycle (*acnB*, *acs*, *fadD*), pyrimidine and deoxyribose degradation (*deoAC*); fatty acid transportation (*fadL*) and fatty acid beta-oxidation by the aerobic (*fadBDE*) and anaerobic (*fadIJ*) pathways. Several anabolic pathways were activated including the fatty acid, lipid and lipopolysaccharide biosynthesis (*accD*, *lpxA*, *rfaD*); peptidoglycan synthesis and recycling (*glmS*, *spr*); synthesis of polyamines (*speB*); nucleotide and histidine biosynthesis (*guaB*, *nrdA*, *thyA*); and porphyrin synthesis from glutamate (*hemL*). Synthesis and degradation of amino acids remained either unaffected or inhibited. Genes encoding multiple ribosomal proteins (*rimM*, *rplI*, *rplL*, *rplU*, *rpsT*, *rpsU*, *rpmA*, *yfgB*) and the protein translation/maturation genes (*efp*, *fklB*) were significantly upregulated that may indicate an increased protein synthesis. The activated anabolism combined with the inhibition of the nutrient uptake systems could lead to a general starvation and catabolic repression by the (p)ppGpp alarmone system that is activated by an increased concentration of uncharged tRNA [30, 31]. To prevent the stringent response, the genes encoding multiple tRNA charging enzymes (*argS*, *glnS*, *glyQ*, *glyS*, *tyrS*) and *spoT* ppGpp hydrolase were upregulated by the treatment with FS-1.

The regulatory network of differential gene expression in the FS-1 treated *E. coli* was modelled by the program PheNetic (Fig. 3). Differentially expressed genes of *E. coli* belonge to several regulons controlled by DNA-binding transcriptional dual regulators CRP, FNR and ArcA, which are involved in transition from aerobic to anaerobic growth; and by three interdependent stress-response sigma-factors, RpoE, RpoS and RpoH, and the osmotic/heavy metal stress response DNA-binding transcriptional regulator CueR. The latter regulator controls the expression of copper transporter CopA and multicopper oxidase CueO, which were significantly upregulated in the FS-1 treated *E. coli*. CRP regulator acts mostly as a transcriptional repressor activated by cAMP in response to the osmotic stress [32]. Also, it was reported that cAMP-CRP complex regulates the central metabolism and carbon source utilization under unavailability of glucose [33].

**Fig. 3.**
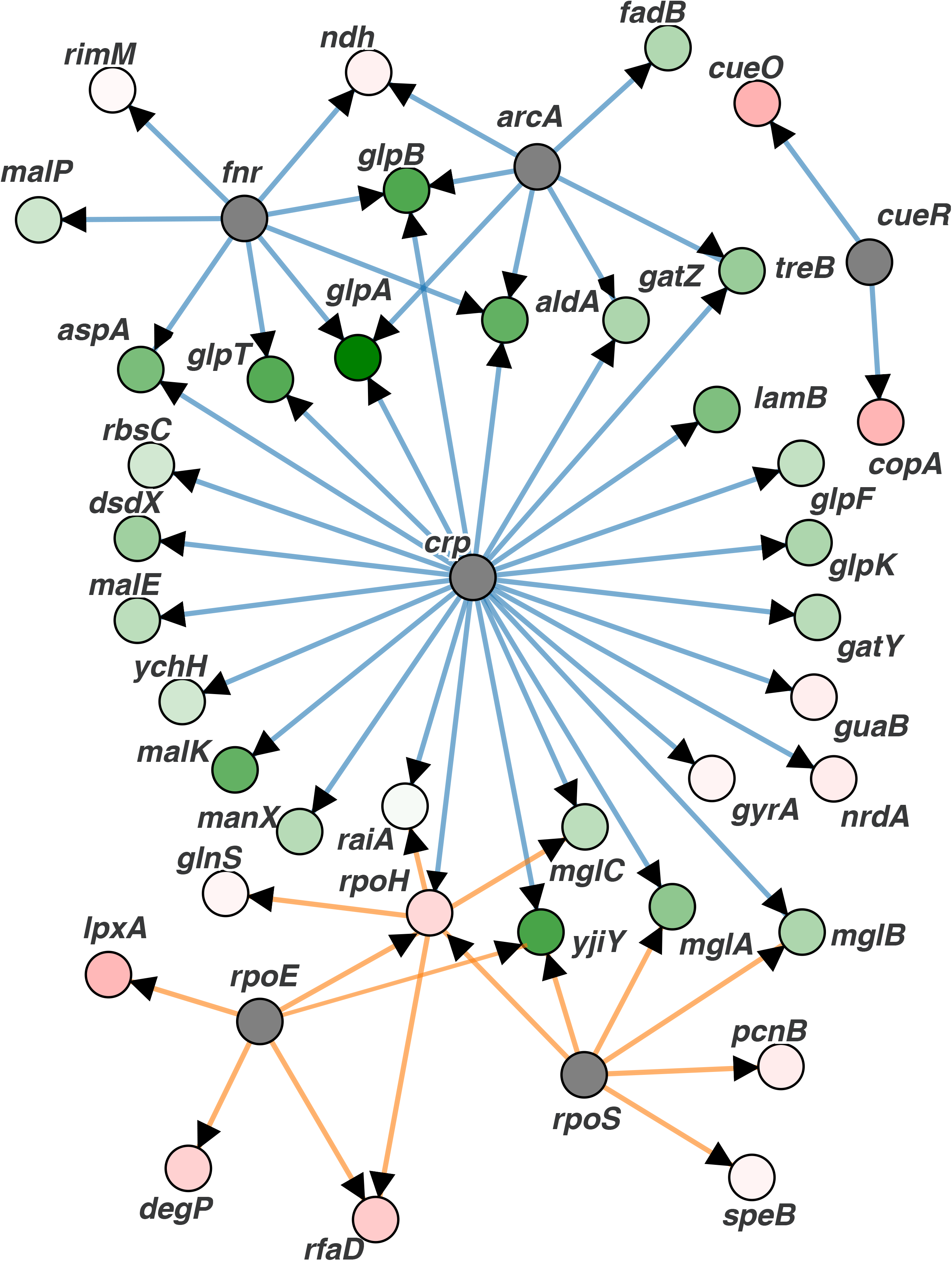
PheNetic network of the regulated genes of *E. coli* BAA-196 identified in the Lag experiment and clustered according to their regulation by higher level transcriptional regulators. Upregulated genes are depicted by pink nodes and downregulated genes – by green nodes (vertices). Colour intensity indicates levels of the expression fold changes. Grey nodes are transcriptional regulators involved in the network, which expression was not reliable changed. Orange edges show regulations by transcriptional activators; and blue edges show regulation by transcriptional repressors. Direct regulations by the transcriptional regulators are indicated by arrowheads.

### Differential gene expression in *S. aureus* ATCC BAA-39

Gene regulation in *S. aureus* treated with FS-1 in the Lag and Log growth phases is shown in Supplementary Table S2. The scheme of the regulated metabolic pathways is shown in Fig. 2B.

The treatment with FS-1 suppressed in *S. aureus* the TCA pathway genes (*icd*, *sucC*) and activated glycolysis (*gpmI*, *pgk*, *tpiA*). This may indicate a possible transition to anaerobiosis [29]; however, the *cydA* gene encoding cytochrome *bd*-I ubiquinol oxidase involved in oxidative phosphorylation process was significantly activated in the FS-1 treated *S. aureus*. Cytochrome *bd*-I terminal oxidase catalyses the reduction of NADH molecules by molecular oxygen or oxygen rich compounds such as nitrates and nitrites. Genes *nirB* and *narGH* encoding nitrite/nitrate reductases involved in dissimilatory nitrite reduction for anaerobic respiration were significantly upregulated in the FS-1 treated *S. aureus* in the Lag growth phase. Another upregulated gene related to anaerobiosis is *fnt* encoding bidirectional formate-nitrite transporter. Fnt plays an important role in regulation of levels of intracellular formate during anaerobic respiration and in providing intracellular dissimilatory nitrite reductases with nitrite. Activation of this transporter may be a preparatory step for anaerobiosis; however, the key enzyme of the anaerobic metabolism, pyruvate:formate lyase PflB that coverts pyruvate to acetyl-CoA and formate, was downregulated in the treated *S. aureus.* Acetyl-CoA synthesis from acetate was activated by an upregulation of acetate kinase *ackA*. However, this enzyme may work in the opposite direction catalysing ATP production from acetyl-CoA. One of the major sources of acetyl-CoA in cells is beta-oxidation of fatty acids. This pathway was inhibited in the FS-1 treated *S. aureus* (Lag-phase experiment) by downregulation of genes *fadA* (2.5-fold) and *fadB* (7.5-fold). Contrary, fatty acid biosynthetic enzyme acetyl-CoA carboxyltransferase *accD* was 1.7-fold upregulated. All these genes were not regulated in the Log-phase experiment. The analysis of the gene expression pattern suggests that *ackA* was activated for acetyl-CoA production. Also, this gene plays an important role for *S. aureus* in maintenance of metabolic homeostasis at stressful conditions [34].

Amino acid biosynthesis was generally downregulated in the FS-1 treated culture except for the glutamate biosynthesis via glutamine (*glnA*, *gltD*) followed by porphyrin biosynthesis (*hemC*) that in *S. aureus* is associated with the growth in anaerobic conditions [35]. Activated synthesis of menaquinone by upregulation of genes *menD* and *menH* also may be associated with anaerobiosis as the encoded compounds are parts of the NADH:menaquinone electron transfer system controlled by NADH oxidoreductase [36].

*De novo* purine (*purA*) and pyrimidine (*pyrG*, *thyA*) biosynthesis and salvage (*guaC*) genes were activated in the Lag-phase experiment as well as the xanthine permease *pbuX* and the hypoxanthine/guanine permease *pbuG*, while the uracil:proton symporter *pyrP* was strongly downregulated. Gene *deoB* associated with nucleotide degradation was downregulated. DNA repair proteins were differentially regulated. Methyl-directed mismatch repair complex *mutSL*, modified pyrimidine reparative endonuclease *nth*, UV DNA repair and Holliday junction renaturation helicases *pcrA* and *ruvB*, RNA decay controlling helicase *cshB*, negative supercoil relaxase *topA*, superoxide dismutase *sodA* and DNA recombination gene *radA* were strongly upregulated. Contrary, SOS-response genes *recA*, *lexA* and *uvrAB* responsible for repair of double-strand DNA breaches were downregulated.

Many genes controlling the accuracy of protein synthesis were activated in the FS-1 treated *S. aureus*. These genes involve tRNA charging (*aspS*, *gltX*, *hisS*, *ileS*, *leuS*, *pheS*, *valS*) and tRNA processing (*gatA*, *gatC*, *miaB*), genes for ribosomal proteins (*rimM*, *rlmB*, *rplJ*, *rpmG*, *rpmH*, *rpsP*, *yjbO*), RNA transcription terminator *rho*, protein release factor *prfC*, protein translocase *secG* and transpeptidase-transglycosylase *sgtA*. Contrary, the ribosomal inhibitor *raiA*, ribosomal methyltransferases *mraW* and *rlmH*, and protein uptake translocase *secA2* were downregulated. Multiple chaperon coding genes, *copZ*, *dnaJ*, *dnaK*, *groL* and *hslO*, were significantly upregulated.

In contrast to *E. coli*, transmembrane transportation proteins generally were not suppressed by the treatment with FS-1 or differentially regulated. The branched chain amino acid transporter *brnQ*, amino acid permease *lysP2*, cysteine transporter *tcyAB* and histidine transporter *yuiF* were upregulated, whereas D-serine/D-alanine/glycine transporter *cycA1*, glutamine transporter *glnQ* and amino acid permease *gabP* were downregulated. Carbohydrate, organic acid and fatty acid transporters *fruA* (fructose), *scrA* (sucrose), *lctP2* (L-lactate) and *msbA* (lipids) were activated, whereas mannitol transporter *mtlA*, oligopeptide transporters *oppA2*, *oppD* and *oppF3*, trehalose transporter *treP* and choline transporters *opuBA* and *opuBB* were inhibited. The multidrug resistance efflux pump *norA*, multiple heavy metal efflux pumps (*copA*, *czrA*, *mntH*, *nikE*, *zitB*), putative sugar efflux transporter HMPRNC0000_0740 and auxin efflux transporter HMPRNC0000_2472 were upregulated.

Transporters of inorganic solutes were significantly activated. These systems include iron and siderophore uptake genes *ceuB* and *sirA*, an operon of ABC iron uptake permeases HMPRNC0000_0811-0813 and ferrichrome binding periplasmic protein HMPRNC0000_2507. Various Na^+^-symporters, *kefC*, *mnhC*, *mnhD*, *nhaC* and HMPRNC0000_0689; as well as K^+^-uptake proteins (*ktrA* and *ntpJ2*) and alkanesulfonate transporter *ssuC* were activated. Potassium transport system Ktr plays an important role in maintenance of the osmotic homeostasis in *S. aureus* [37]. Other osmoprotective compounds, glycine betaine and polyamines, were actively synthesized (*betB*) or transported to the cell (*opuD2*, *potA*, *potC*). Osmotic stress may occur due to damaging of the bacterial cell walls by iodine. Genes involved in cell wall biosynthesis were regulated differentially. The peptidoglycan synthesis, salvage and stress response genes, N-acetylmuramoyl-L-alanine amidase *atl*, peptidoglycan DD-endopeptidase *dacB*, cell wall stress response autolysin *amiD2* and cell wall biosynthesis genes *fmtB2*, *murD* and *murE* were upregulated. Contrary, D-alanylalanine ligase *ddl*, peptidoglycan transglycosylase *mgt*, N-acetylmuramic acid 6-phosphate etherase *murQ* and N-acetylmuramic acid-specific transporter *sacX* were downregulated.

### Differential gene expression in *A. baumannii* ATCC BAA-1790

Gene regulation in *A. baumannii* treated with FS-1 in the Lag and Log growth phases is shown in Supplementary Table S3. The scheme of the regulated metabolic pathways is shown in Fig. 2C.

Exposure of *A. baumannii* to FS-1 has activated the transcription of citrate synthase *gltA-2* and isocitrate lyase *aceA* involved in the TCA and glyoxylate cycle. These pathways were strongly inhibited by FS-1 in *E. coli* and *S. aureus*. Upregulation of phosphoenolpyruvate carboxykinase *pck* indicated an activation of the gluconeogenetic pathway, whereas the glycolysis was not activated by FS-1 in contrast to the two former model microorganisms. The activation of the anabolic gluconeogenetic pathway may be associated with the intensified synthesis of various amino acids observed in the FS-1 treated *A. baumannii*. Genes associated with the synthesis of branched chain amino acids (*ilvC*, *ilvH*, *ilvI*, *leuB*, *leuC*, *leuD*), aromatic amino acid (*aroF*), cysteine (*cysD*, *cysI*, *cysN*), arginine (*argG*), glutamate (*gltD*), methionine, lysine and threonine (*metE*, *metB*, *usg*), and S-adenosylmethionine (*metK*, *mmuM*) were upregulated at least in one or both growth phases. These pathways were generally inhibited in the FS-1 treated *E. coli* and *S. aureus*.

Cytochrome *bd*-I terminal oxidase CydAB showing an upregulation in the FS-1 treated *E. coli* and *S. aureus*, was suppressed in the FS-1 treated *A. baumannii*. In the text above it was hypothesized that the activation of this cytochrome complex together with the TCA downregulation and the activation of glycolysis might be indicative for the transition of bacteria to anaerobic respiration that possibly was not the case with the FS-1 treated *A. baumannii*. The aerobic fatty acid oxidation pathway was upregulated (*fadAB*) while the anaerobic branch of this pathway (*fadK*) was downregulated.

Genes involved in arginine catabolism to ammonia (*astA*, *astB*, *astD*, *astE*) were downregulated that opposed the reaction of FS-1 treatment in *E. coli* and *S. aureus*. The treatment of *A. baumannii* with FS-1 has activated ribosomal inhibitors *rsfA* and *yfiA* while the pattern of upregulated and downregulated ribosomal proteins was dissimilar to that in the FS-1 treated *E. coli* and *S. aureus* with many of these genes being strongly downregulated. Genes *pyrH*, *lpxH* and *fabF* involved in the nucleotide and fatty acid biosynthesis were downregulated while the nucleotide degradation and salvage genes *ppnP* and *rutF* were upregulated. A strong downregulation was observed for the potassium uptake proteins KdpABCD, which play an important role in maintenance of osmotic homeostasis [38, 40] and in the adaptation of *E. coli* to the presence of FS-1 in the medium [26]. Peptidoglycan synthesis and assembly genes *murI* and *dacA*; and heavy metal resistance genes *copA* and *spxA* were strongly upregulated in FS-1 treated *A. baumannii* as it was observed also in the FS-1 treated *E. coli* and *S. aureus*.

The activation of many genes involved in the thioredoxin pathway, glutathione-glutaredoxin redox reactions and oxidative stress response (*gntY*, *nfnB*, *usg*, *nemA*, *osmC* and *wrbA-2*) suggests that the FS-1 treated culture of *A. baumannii* experienced an oxidative stress. In the FS-1 treated *E. coli* and *S. aureus*, the homologous genes were not regulated, while other genes controlling the redox homeostasis showed some levels of downregulation: *grxA* and *trxC* in *E. coli*; *gpo* and *osmC* in *S. aureus*.

Multiple genes involved in iron scavenging and uptake were activated in the FS-1 treated *A. baumannii*, while gene *bfr-1* for iron storage in bacterioferritin was downregulated. The sulphate and thiosulphate uptake and assimilation genes (*cysA*, *cysB*, *cysP*, *cysW* and *cysU*) were strongly upregulated in the Log-phase that probably was associated with the activation of biosynthesis of sulphate-containing amino acids cysteine, methionine and S-adenosylmethionine. The same was true for the observed upregulation of *tauAB* controlling the uptake and assimilation of sulphate-rich compound taurine. Nitrate/sulfonate/bicarbonate ABC transporter *ybaN* collocated on the chromosome with *tauAB* operon was also activated. In the Lag-phase, all these genes were strongly downregulated. Contrary, phosphate ABC transporter permease subunits *pstA* and *pstB* were downregulated by the treatment with FS-1 in the Log-phase but upregulated in the Lag-phase. Several other genes showed an opposite regulation in the Lag and Log experiments on *A. baumannii* including sigma factor *rpoH* and transcriptional regulator *merR*, which were upregulated by the treatment with FS-1 in the Lag-phase but downregulated in the Log-phase. Differential expression of these top-level transcriptional regulators may be behind the opposite regulation of other above-mentioned genes in the Lag and Log experiments.

### Transcriptional regulation of the genes located on the plasmids of *E. coli* ATCC BAA-196 and *A. baumannii* ATCC BAA-1790

Genomes of the two model microorganisms used in this study, *E coli* BAA-196 and *A. baumannii* BAA1790, comprise the plasmids of 266,396 bp and 67,023 bp, respectively. There are 264 predicted protein coding genes on the *E. coli* plasmid, but almost all of them were transcriptionally silent at the experimental and the negative control conditions. Low level transcription was recorded only for 8 transposases, two beta-lactamases and two aminoglycosidase acetyltransferases. The treatment of these bacteria with FS-1 in the Lag-phase suppressed the transcription from all these genes and they showed a differential regulation in the Log growth phase. However, the observed changes in the levels of gene expression in both experiments were statistically unreliable with *p*-values much higher than 0.05. A previously published study, where the strain *E. coli* BAA-196 was cultivated with FS-1 for 10 passages, showed a significant expression from almost all the genes located on the plasmid suggesting its importance for protection of this bacterium from environmental stresses [27]. Possibly, the five-minutes treatment with FS-1 was insufficient for the plasmid activation.

The *A. baumannii* plasmid comprises 82 protein coding genes. Almost all these genes showed some levels of expression when the strain was cultivated on the medium without FS-1. Many of these genes were differentially regulated by FS-1. The highest level of up- and downregulation was observed for the genes encoding plasmid conjugation proteins. However, in all these cases the observed differential gene expression was statistically unreliable due to significant variations in the levels of expression of these genes in repeated experiments.

### Comparison of gene expression profiles of three model microorganisms treated with FS-1

To compare the patterns of gene expression, the program GET-HOMOLOGOUS [38] was used to identify homologous genes shared by the model microorganisms. A summary of this search is represented by Venn diagram in Supplementary Fig. S2. There were 504 genes shared by the three genomes and additionally 839 genes were common for *E. coli* and *A. baumannii*; 221 homologs were shared by *E. coli* and *S. aureus*; and 122 homologous genes were shared by *S. aureus* and *A. baumannii*.

Plots of gene expression fold changes determined for the homologous genes in pairs of model organisms, A) *E. coli* and *S. aureus*; B) *E. coli* and *A. baumannii*; C) *S. aureus* and *A. baumannii*; all treated with FS-1 in the Lag-phase, are shown in corresponding parts of Fig. 4. Calculated Pearson correlation coefficients were in the range from 0.281 for the pair *E. coli* ─ *S. aureus* (Fig. 4A) to 0.007 for the pair *S. aureus* ─ *A. baumannii* (Fig. 4C). Only one gene was found to show a significant upregulation in all three tested microorganisms: heavy metal (copper, lead, cadmium, zinc and mercury) transporter *copA*. It suggests that the treatment with FS-1 has increased penetrability of the cells for heavy metal ions in all three tested bacteria probably due to damaging cell wall and cytoplasmic membrane barrier molecules by iodine. In response to the influx of heavy metal ions, the corresponding efflux pumps were activated. This finding agrees with another study reporting a disintegration of bacterial membranes by iodine nanoparticles [18]. Other authors reported that the copper homeostasis systems may be activated in *E. coli* in anaerobic conditions associated with amino acid limitation [41].

**Fig. 4.**
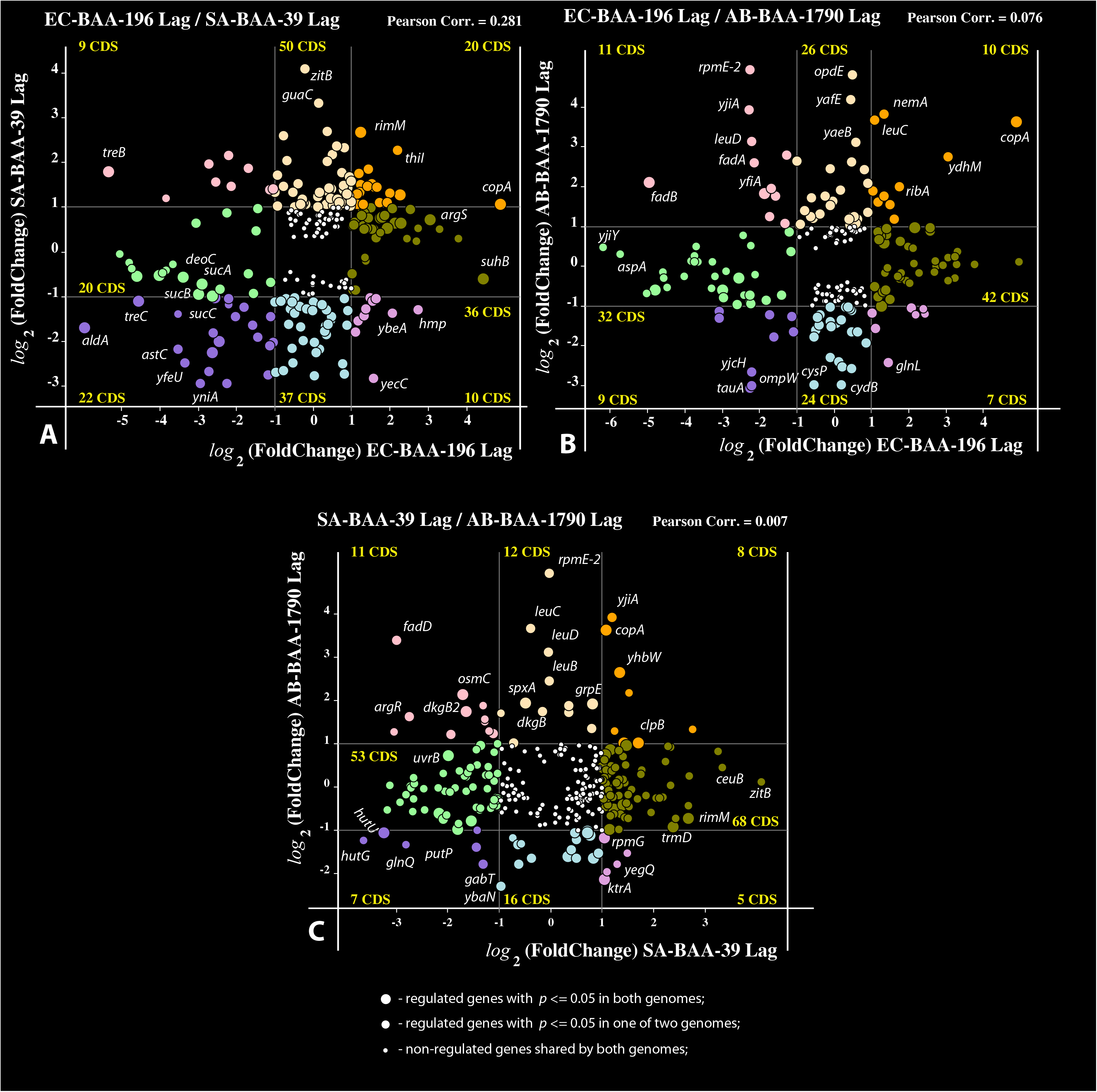
Plots of co-regulation of homologous gene in the Lag experiment shared by pairs of model microorganisms. A) *E. coli* BAA-196 and *S. aureus* BAA-39 (EC-BAA-196 / SA-BAA-39); B) *E. coli* BAA-196 and *A. baumannii* BAA-1790 (EC-BAA-196 /AB_BAA-1790); and C) *S. aureus* BAA-39 and *A. baumannii* BAA-1790 (SA-BAA-39 / AB-BAA-1790). Circles represent protein coding genes (CDS) plotted according to their negative and positive *log*_2_(fold-change) values calculated in the Lag-experiments for different microorganisms shown along axes X and Y. The outermost regulated genes are labelled by their names. Thin vertical and horizontal lines within the plots separate genes with 2-fold or higher regulation and split the plots to sectors of genes of different categories depending on their coregulation. Numbers of CDS falling to different sectors are shown. Up- and down-coregulated genes and oppositely regulated genes are depicted by different colours. Statistical reliability of the fold change predictions is depicted by sizes of the circles as explained in the legend in the bottom part of the figure. Estimated Pearson correlation coefficients are given on the top of each plot.

The close-to-zero level of gene coregulation in these three phylogenetically distant bacteria treated with FS-1 may be explained by the fact that the regulation of similar metabolic processes often is carried out by up- and downregulation of different genes involved in the metabolic pathways. Indeed, the comparison of the affected metabolic pathways of all these bacteria (Table 3 and Fig. 2) demonstrated a higher similarity in their responses to the treatment with FS-1. This commonality includes downregulation of many transmembrane transporters of carbohydrates, amino acids, oligopeptides and oligosaccharides in all these bacteria. FS-1 is a complex of iodine molecules bound to dextrin fragments and oligopeptides. Transporters of these compounds may serve as entrance points for iodine to the cells. This may explain the observed inhibition of these transporters; however, the spectra of inhibited and activated transporters were different in the tested bacterial strains. Sodium-proline symporter *putP* was downregulated in all three tested bacteria almost at all conditions (except for *E. coli* in the Log-phase). Iron scavenging and uptake proteins were activated in *S. aureus* and *A. baumannii*, probably due to immobilizing of iron ions by iodine into insoluble iodides; however, in *E. coli* the expression of these genes was not regulated. Other systems of transmembrane uptake and utilization of various compounds were regulated differentially in the tested bacteria.

Synthesis of cell wall compounds including peptidoglycans and lipids was generally upregulated. This observation confirms that the prime target of the released iodine was the cell wall and cell membrane structures that agrees with other studies [18, 21]. Various efflux pumps including several heavy metal resistance proteins were activated by the treatment with FS-1. The activation of synthesis and uptake of osmoprotective molecules (glycine betaine and polyamines) and the potassium uptake systems in *E. coli* and *S. aureus* suggests that these strains experienced an osmotic stress caused by the treatment with FS-1. In *E. coli*, an important role in gene regulation under the experimental condition was played by cAMP-CRP signal transduction system (Fig. 3); however, it was not the case with the gene regulation in the FS-1 treated *S. aureus* and *A. baumannii*, which probably did not utilize cAMP-CRP for gene expression regulation. Particularly, adenylate cyclase *cyaA* was strongly downregulated in the FS-1 treated *S. aureus*.

The effect of FS-1 on *A. baumannii* in the Log growth phase was rather specific and characterized with a significant downregulation of many genes including several important transcriptional regulators: EnvZ/OmrR two-component signal transduction system involved in osmoregulation [42], RstA response regulator of many intracellular processes [43], potassium-dependent KdpD regulator controlling K^+^-homeostasis [44], iron uptake regulator *fur*, sigma factor-32 *rpoH* and transcriptional regulator *merR*.

## Discussion

Noteworthy finding of this work was that the tested model microorganisms responded differently to the short-time (5 min) treatment with the iodine-containing nano-micelles. The treatment of *E. coli* and *S. aureus* with FS-1 caused a transition of these bacteria to anaerobiosis. One possible reason for this reaction may consist in damaging the O_2_ dependent electron transportation systems by I_2_ molecules. This caused bacteria to look for alternative electron acceptors such as DMSO in the case with *E. coli* or nitrates/nitrites in the case with *S. aureus*. In another study on *E. coli* BAA-196 cultivated for 10 passages with FS-1, a complete stall of expression of genes encoding cytochrome *bo* terminal oxidase complex (*cyoA*, *cyoB*, *cyoC* and *cyoE*) was observed [26, 28] that corroborates with the hypothesis that iodine may interfere with the systems of electron transportation between cytochromes and oxygen. This transition to anaerobiosis caused by the treatment with FS-1 may be of practical importance. Both these bacteria, *E. coli* and *S. aureus*, may survive the anaerobic or microaerophilic conditions; however, the transfer to anaerobiosis occurs at the expense of reduction of their virulence and antibiotic resistance [45].

Another reason for the transition to anaerobiosis may consist in an attempt to alleviate the oxidative stress caused by iodine in the medium as the oxidative phosphorylation is a source of peroxide radicals aggravating the oxidative stress. This hypothesis is supported by the observed strong oxidative stress response in the FS-1 treated *A. baumannii*, which in contrast to *E. coli* and *S. aureus* did not show any signs of transition to anaerobiosis. Several other incongruent responses by *A. baumannii* to the treatment with FS-1 compared to other two microorganisms, such as the activation of gluconeogenesis over glycolysis and the enzymes involved in TCA, glyoxylate pathway, fatty acid beta-oxidation and amino acid biosynthesis with opposite effects in the FS-1 treated *E. coli* and *S. aureus* may be attributed to the transition of the latter strains to anaerobiosis with the absence of such a response in *A. baumannii*. The gene expression profiles in *A. baumannii* under the effect of FS-1 differed in the Lag- and Log-phases with several genes involved in sulphur, nitrogen and phosphorus uptake being regulated oppositely in these two experiments.

The treatment of bacteria with the iodine-containing micelle drug FS-1 caused the osmotic and oxidative stresses to the bacterial cells. The osmotic stress was associated with damaging of cell wall and cytoplasmic membrane proteins leading to an increased penetrability of bacterial cells to heavy metal ions and possibly to antibiotics. This effect was more apparent to *E. coli* and *S. aureus*. The oxidative stress could be partly alleviated in the FS-1 treated *E. coli* and *S. aureus* due to the transition to anaerobiosis. However, the strong inhibition of multiple nutrient uptake-and-utilization systems in *E. coli* could lead this bacterium to a shortage of energy and nutrients. In *S. aureus*, the elevated accumulation of iodine in cytoplasm caused stronger damaging of DNA that was apparent by the observed activation of DNA repair and nucleotide salvage systems. Another study when *S. aureus* was cultivated for long time with FS-1 showed an increased rate of mutations and abnormal epigenetic modifications of the chromosomal DNA [26]. This activation of the DNA repair systems in response to the treatment with FS-1 was not observed in other two bacteria. Cell wall of *A. baumannii* was apparently the most resistant against iodine and prevented cells from iodine penetration into cytoplasm. However, this bacterium was under a stronger oxidative stress compared to the other two microorganisms.

## Conclusion

The three tested pathogenic bacteria responded differently to the treatment with the iodine-containing nano-micelles depending also on the growth phase when the drug was applied. One common mechanism was the activation of the efflux pump CopA that probably points on damaging cell wall structures in the FS-1 treated bacteria leading to an increased permeability of the cytoplasmic membrane and the osmotic stress. This information will be of importance for development of new iodine-containing nano-micelle drugs against antibiotics resistant nosocomial infections represented in this study by the reference multidrug resistant strains, *E. coli* ATCC BAA-196, *S. aureus* ATCC BAA-39 and *A. baumannii* ATCC BAA-1790.

Further studies will aim at testing the effects of iodine-containing compounds on a broader collection of antibiotic resistant nosocomial infection agents isolated from clinics in several recent studies [46, 47]. Although iodine is in the use by humankind as an antimicrobial disinfectant for more than two centuries, to our best knowledge till now there have been no published reports on the effect of iodine applied in sublethal concentrations in nano-micelle complexes on the gene expression and metabolism of pathogenic bacteria.

## Materials and methods

### Bacterial cultures

Multidrug resistant strains *Staphylococcus aureus* subsp. *aureus* ATCC^®^ BAA-39™, *Escherichia coli* ATCC^®^ BAA-196*™* and *Acinetobacter baumannii* ATCC^®^ BAA-1790*™* were obtained from the American Type Culture Collection (ATCC) and used as the model organisms in this study. Stock cultures were kept in a freezer at −80°C. Bacteria were cultivated on Mueller-Hinton (MH) liquid or solid media (Himedia, India) without antibiotics (*S. aureus* and *A. baumannii*) and in the medium supplemented with ceftazidime 10 μg/ml (*E. coli*) as recommended in the ATCC product sheets. Before the use in the experiments, the strains were twice passaged on MH medium.

### Determination of growth rates

Overnight (24 hours) culture growth was adjusted by dilution with sterile water to 1 × 10^7^ CFU/ml and inoculated into microarray plates with MH liquid medium. Plates were incubated at 37°C on 60 rpm shaker for 24 hours with optical density (OD at 620 nm) recording every 5 min by Multiscan Ascent spectrophotometer (Thermo, Finland). The growth curves were estimated based on the obtained series of OD values in three repeats by Baranyi and Roberts model implemented in the interactive DMFit application [48] as shown in Supplementary Fig. S1.

### Treatment with FS-1

Bacterial inoculants were incubated at 37°C as follows: *E. coli* for 1 hour (the end of the lagging growth phase, termed Lag-experiment) and for 6 hours (the mid of the logarithmic growth phase, Log-experiment); *S. aureus* for 2.5 hours (Lag) and 9 hours (Log); and *A. baumannii* for 1 hour (Lag) and 10 hours (Log). Lag and Log growth phases were estimated based on the strain specific growth curves on MH medium (Supplementary Fig. S1). Thereafter, the experimental cultures were suplemented with FS-1 to achieve the final concentration of 450 μg/ml (for *S. aureus*) or 500 μg/ml (for *E. coli* and *A. baumannii*) that corresponded to the half of the minimal bactericidal concentrations of FS-1 estimated for these bacteria; whereas for the negative control samples, the same volume of physiological saline was added to the bacterial cultures. Cultures were incubated for 5 min at 37°C with shaking. Metabolic processes were stopped by adding the killing buffer (2.0 ml of 1 M Tris-HCl, pH 7.5; 0.5 ml of 1 M MgCl_2_, 1.3 g of NaN_3_; 997.5 ml of water) in the volume ratio 1:1 [49]. Bacterial cells were collected for RNA extraction by centrifugation at 5,000 g for 10 min.

### RNA Library Preparation and Sequencing

Total RNA samples were extracted from the cultures with the use of the RiboPure Bacteria Kit (Ambion, Lithuania) as instructed by the developer’s guidelines. Afterwards, the quantity and quality of the extracted RNA was verified with the use of the NanoDrop 2000c spectrophotometer (Thermo Scientific, USA) at optical wavelengths 260 and 280 nm. Purification of the total RNA from ribosomal 16S and 23S RNA was carried out using the MICROBExpress Bacterial mRNA Purification Kit (Ambion, Lithuania) following the developer recommendations. Thereafter, the effectiveness of sample purification was determined on the Bioanalyzer 2100 (Agilent, Germany) with the RNA 6000 Nano LabChip Kit (Agilent Technologies, Lithuania).

The library preparation from the extracted RNA samples included an enzymatic fragmentation step with the use of the Ion Total RNA Seq kit V2. Thereafter, the Ion Xpress RNA-Seq Barcode 01-16 Kit was used for RNA fragment barcoding. cDNA sequencing was then performed using the Ion 318 Chip Kit V2 on the Ion Torrent PGM sequencer (Life Technologies, USA).

### Differential gene expression analyses

Gene expression in bacterial cells treated with FS-1 for 5 min was compared to gene expression profiles in bacterial cells treated with the same volume of saline (negative control). RNA extraction from bacterial cells treated with FS-1 and grown at the negative control condition was repeated three times with *E. coli* BAA-196 and six times with *S. aureus* BAA-39 and *A. baumannii* BAA-1790. RNA samples were collected from different test-tubes with bacterial growth incubated in parallel to ensure biological rather than technical replication of the results.

Sampled RNA was converted to cDNA and sequenced as explained above. Generated DNA reads were quality trimmed and filtered using the raw DNA read processing pipeline implemented in Unipro UGENE v1.32.0 [50] using 20 as the quality threshold. Reads shorter than 30 bp were filtered out.

The differential gene expression was estimated using the R-3.4.4 software packages. Firstly, reference indices were built for each reference genome using the *buildindex* function available in the *Rsubreads* package (Bioconducter, www.bioconductor.org). For each bacterium, the obtained RNA fragments were aligned to the relevant reference genomes in FASTA formats (*S. aureus* ATCC BAA-39 NC [CP033505.1], *E. coli* ATCC BAA-196 NC [CP042865.1 for chromosome and CP042866.1 for plasmid], *A. baumannii* ATCC BAA-1790 [CP042841.1 for chromosome and CP042842.1 for plasmid]). Resulted read alignment files in BAM format were sorted using Samtools software package available from htslib (www.htslib.org) and used for counting of reads overlapping predicted gene locations by using the *featureCounts* function and the relevant genome annotation files in GFF format. The R packages *DESeq2* (Bioconducter) and *GenomicFeatures* tools were then used for normalization of read counts and calculation of the differential gene expression statistics. All the above-mentioned tools were pipelined using an in-house Python 2.7 script available from the SeqWord project Web-site http://seqword.bi.up.ac.za/transcriptomics_scripts/. This script also was used for visualization of the results as shown in Fig. 1 and 4.

The numbers of quality-controlled DNA reads generated in different experiments and the numbers of reads uniquely mapped against CDS and rRNA sequences of the reference bacterial genomes are summarized in Table 1.

### Metabolic pathway and regulatory network analyses

With the use of identified differentially expressed genes, metabolic pathways and reactions controlled by those genes were identified using the Pathway Tools v24.0 software [51]. Networks of regulated genes were constructed using the Web based tool PheNetic [52] based on the regulation network designed for *E. coli* K12 [NC_000913. 2] and available from the PheNetic Web site. Pairs of homologous genes in the genomes K12 and BAA-196, and also the homologous genes shared by *S. aureus* BAA-39, *E. coli* BAA-196 and *A. baumannii* BAA-1790 were identified using the program GET_HOMOLOGUES with the parameters set by default [38]. Large admixed clusters of homologous phage-related integrases and transposases were considered as strain-specific genes and excluded from the transcription comparison. Venn diagram with numbers of shared genes is shown in Supplementary Fig. S2.

### Statistical evaluations

All RNA sequencing experiments were performed in six (*S. aureus* and *A. baumannii*) or three (*E. coli*) repetitions. Genes showing 2-fold or greater expression difference with calculated *p*-value equal or smaller 0.05 were considered as significantly regulated. Aggregated Pearson correlation coefficients (*C*_*p*_) of gene coregulation were calculated by Eq. 1 implemented in an in-house Python 2.7 script mentioned above.

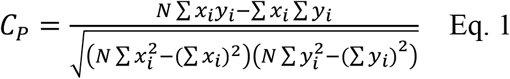

where *x*_*i*_ and *y*_*i*_ are fold-change values estimated for every *i*’s homologous gene, *N* – total number of shared homologous genes.

## Data availability

Experimental and negative control RNA reads generated in this study are freely available from BioProject Web-sites of the reference genomes deposited at NCBI: *S. aureus* BAA-39 NC, PRJNA480363; *E. coli* BAA-196 NC, PRJNA557356; *A. baumannii* BAA-1790 NC, PRJNA557366.

## Acknowledgements

Authors of the paper wish to thank the head of the Microbiology department of the Scientific Center for Anti-Infectious Drugs (Alamty, Kazakhstan), Dr. Ardak B. Jumagaziyeva, and the staff members of the Microbiology department for maintaining the model microorganisms received from the ATCC and keeping them readily available for this study.

---

Supplementary Table S1. Genes regulated in *E. coli* ATCC BAA-196 in Lag and Log growth phases by 5 min exposure to FS-1.

Supplementary Table S2. Genes regulated in *S. aureus* ATCC BAA-39 in Lag and Log growth phases by 5 min exposure to FS-1.

Supplementary Table S3. Genes regulated in *A. baumannii* ATCC BAA-1790 in Lag and Log growth phases by 5 min exposure to FS-1.

**Supplementary Fig. S1.** Growth curves of A) *E. coli* ATCC BAA-196; B) *S. aureus* ATCC BAA-39; C) *A. baumannii* ATCC BAA-1790 calculated by Baranyi and Roberts model implemented in the interactive DMFit application based on optical density (OD) values obtained in three repetitions using microarray plates with MH liquid medium. Average OD values are depicted by open circles. R-square values of deviations of predicted from recorded OD values are shown.

**Supplementary Fig. S2.** Venn diagram of distribution of strain-specific and homologous core genes shared by *E. coli* ATCC BAA-196, *S. aureus* ATCC BAA-39 and *A. baumannii* ATCC BAA-1790 as predicted by the program GET_HOMOLOGUES. Large admixed clusters of homologous phage-related integrases and transposases were considered as strain-specific genes.

## References

1. Kelly FC. 1961. Iodine in Medicine and Pharmacy Sinceits Discovery – 1811-1961. Proc R Soc Med 54:831–836.

2. Abraham EP, Chain E, Fletcher CM, Florey HW, Gardner AD, Heatley NG, Jennings MA. 1992. Further observations on penicillin. 1941. Eur J Clin Pharmacol 42:3–9.

3. Munita JM, Arias CA. 2016. Mechanisms of Antibiotic Resistance. Microbiol Spectr 4:10.1128/microbiolspec.VMBF-0016-2015. doi:10.1128/microbiolspec.VMBF-0016-2015.

4. Barbosa TM, Levy SB. 2000. The impact of antibiotic use on resistance development and persistence. Drug Resist Updat 3:303–311. doi:10.1054/drup.2000.0167.

5. Nicolaou KC, Montagnon T. 2008. Molecules that changed the world. Weinheim, Wiley-VCH.

6. Zucca M, Savioa D. 2010. The Post-Antibiotic Era: promising developments in the therapy of infectious diseases. Int J Biomed Sci 6:77–86.

7. Laxminarayan R. 2014. Antibiotic effectiveness: balancing conservation against innovation. Science 345:1299–1301. doi:10.1126/science.1254163.

8. Ventola CL. 2015. The antibiotic resistance crisis: part 1: causes and threats. P T 40:277–283.

9. Bush K, Courvalin P, Dantas G, Davies J, Eisenstein B, Huovinen P, Jacoby GA, Kishony R, Kreiswirth BN, Kutter E, Lerner SA, Levy S, Lewis K, Lomovskaya O, Miller JH, Mobashery S, Piddock LJ, Projan S, Thomas CM, Tomasz A, Tulkens PM, Walsh TR, Watson JD, Witkowski J, Witte W, Wright G, Yeh P, Zgurskaya HI. 2011. Tackling antibiotic resistance. Nat Rev Microbiol 9:894–896. doi:10.1038/nrmicro2693.

10. McClure NS, Day T. 2014. A theoretical examination of the relative importance of evolution management and drug development for managing resistance. Proc Biol Sci 281:20141861. doi:10.1098/rspb.2014.1861.

11. Anguiano B, García-Solís P, Delgado G, Aceves Velasco C. 2007. Uptake and gene expression with antitumoral doses of iodine in thyroid and mammary gland: evidence that chronic administration has no harmful effects. Thyroid 17:851–859. doi:10.1089/thy.2007.0122.

12. Stoddard FR 2nd, Brooks AD, Eskin BA, Johannes GJ. 2008. Iodine alters gene expression in the MCF7 breast cancer cell line: evidence for an anti-estrogen effect of iodine. Int J Med Sci 5:189–196. doi: 10.7150/ijms.5.189.

13. Kitagawa E, Akama K, Iwahashi H. (2005). Effects of iodine on global gene expression in *Saccharomyces cerevisiae*. Biosci Biotechnol Biochem 69:2285–2293. doi:10.1271/bbb.69.2285.

14. Klebanoff SJ. 1967. Iodination of bacteria: a bacteriocidal mechanism. J Exp Med 126:1063–1078. PMID: 4964565.

15. Pogue JM, Marchaim D, Kaye D, Kaye KS. 2011. Revisiting “older” antimicrobials in the era of multidrug resistance. Pharmacotherapy 31:912–921. doi:10.1592/phco.31.9.912.

16. Banerjee M, Mallick S, Paul A, Chattopadhyay A, Ghosh SS. 2010. Heightened reactive oxygen species generation in the antimicrobial activity of a three component iodinated chitosan-silver nanoparticle composite. Langmuir 26:5901–5908. doi:10.1021/la9038528.

17. Murdoch R, Lagan KM. 2013. The role of povidone and cadexomer iodine in the management of acute and chronic wounds. Physical Therapy Reviews 18:207–216. doi:10.1179/1743288X13Y.0000000082.

18. Viswanathan K, Babu DB, Jayakumar G, Raj GD. 2017. Anti-microbial and skin wound dressing application of molecular iodine nanoparticles. Materials Research Express 4:104003.

19. Dandia A, Gupta SL, Maheshwari S. 2014. Molecular iodine: mild, green, and nontoxic lewis acid catalyst for the synthesis of heterocyclic compounds, p. 277–327. *In* Ameta KL, Dandia A. (eds.), Green Chemistry: Synthesis of Bioactive Heterocycles. Springer, New Delhi, India.

20. Moulay S. 2013. Molecular iodine/polymer complexes. J Polymer Eng 33:389–443.

21. Ilin A, Kerimzhanova B, Yuldasheva G. 2017. Action mechanism of molecular iodine complex with bioorganic ligands, magnesium and lithium halogenides on human tuberculosis strain with multiple drug resistance. Journal of Microbial and Biochemical Technology 9:293–300.

22. Ilin AI, Kulmanov ME, Korotetskiy IS, Islamov RA, Akhmetova GK, Lankina MV, Reva ON. 2017. Genomic insight into mechanisms of reversion of antibiotic resistance in multidrug resistant *Mycobacterium tuberculosis* induced by a nanomolecular iodine-containing complex FS-1. Front Cell Infect Microbiol 7:151. doi:10.3389/fcimb.2017.00151.

23. Joubert M, Reva ON, Korotetskiy IS, Shvidko SV, Shilov SV, Jumagaziyeva AB, Kenesheva ST, Suldina NA, Ilin AI. 2019. Assembly of complete genome sequences of negative-control and experimental strain variants of *Staphylococcus aureus* ATCC BAA-39 selected under the effect of the drug FS-1, which induces antibiotic resistance reversion. Microbiol Resour Announc 8:e00579–19. https://doi.org/10.1128/MRA.00579-19.

24. Korotetskiy IS, Joubert M, Taukobong S, Jumagaziyeva AB, Shilov SV, Shvidko SV, Suldina NA, Kenesheva ST, Yssel A, Reva ON, Ilin AI. 2019. Complete genome sequence of a multidrug-resistant strain, *Escherichia coli* ATCC BAA-196, as a model for studying induced antibiotic resistance reversion. Microbiol Resour Announc 8:e01118–19. doi:10.1128/MRA.01118-19.

25. Korotetskiy IS, Joubert M, Magabotha SM, Jumagaziyeva AB, Shilov SV, Suldina NA, Kenesheva ST, Yssel A, Reva ON, Ilin AI. 2020. Complete genome sequence of a collection strain *Acinetobacter baumannii* ATCC BAA-1790 used as a model to study the antibiotic resistance reversion induced by the iodine-containing complexes. Microbiol Resour Announc 9:e01467–19. doi: 10.1128/MRA.01467-19.

26. Reva ON, Korotetskiy IS, Joubert M, Shilov SV, Jumagaziyeva AB, Suldina NA, Ilin AI. 2020.The effect of iodine-containing nano-micelles, FS-1, on antibiotic resistance, gene expression and epigenetic modifications in the genome of multidrug resistant MRSA strain *Staphylococcus aureus* ATCC BAA-39. Front. Microbiol. 11. doi: 10.3389/fmicb.2020.581660.

27. Korotetskiy IS, Jumagaziyeva AB, Shilov SV, Kuznetsova TV, Suldina NA, Kenesheva ST, Ilin AI, Joubert M, Taukobong S, Reva ON. 2020. Differential gene expression and alternation of patterns of DNA methylation in the multidrug resistant strain *Escherichia coli* ATCC BAA-196 caused by iodine-containing nano-micelle drug FS-1 that induces antibiotic resistance reversion. bioRxiv. doi: 10.1101/2020.05.15.097816.

28. Korotetskiy IS, Shilov SV, Reva ON, Kuznetsova TV, Jumagaziyeva AB, Akhmatullina NB, Ilin AI. 2020. Gene expression profiling of multi-drug resistant *E. coli* after exposure by nanomolecular iodine-containing complex. News Nat Acad Sci of the Republic of Kazakhstan 4:2224–5308. doi: 10.32014/2020.2519-1629.27.

29. Kohler C, von Eiff C, Liebeke M, McNamara PJ, Lalk M, Proctor RA, Hecker M, Engelmann S. 2008. A defect in menadione biosynthesis induces global changes in gene expression in *Staphylococcus aureus*. J Bacteriol 190:6351–6364. doi:10.1128/JB.00505-08.

30. Rojiani MV, Jakubowski H, Goldman E. 1989. Effect of variation of charged and uncharged tRNA(Trp) levels on ppGpp synthesis in *Escherichia coli*. J Bacteriol 171:6493–6502. doi:10.1128/jb.171.12.6493-6502.1989.

31. Chatterji D, Ojha AK. 2001. Revisiting the stringent response, ppGpp and starvation signaling. Curr Opin Microbiol 4:160–165. doi:10.1016/s1369-5274(00)00182-x.

32. Balsalobre C, Johansson J, Uhlin BE. 2006. Cyclic AMP-dependent osmoregulation of *crp* gene expression in *Escherichia coli*. J Bacteriol 188:5935–5944. doi: 10.1128/JB.00235-06.

33. Shimada T, Fujita N, Yamamoto K, Ishihama A. 2011. Novel roles of cAMP receptor protein (CRP) in regulation of transport and metabolism of carbon sources. PLoS One 6:e20081. doi: 10.1371/journal.pone.0020081.

34. Sadykov MR, Thomas VC, Marshall DD, Wenstrom CJ, Moormeier DE, Widhelm TJ, Nuxoll AS, Powers R, Bayles KW. 2013. Inactivation of the Pta-AckA pathway causes cell death in *Staphylococcus aureus*. J Bacteriol 195:3035–3044. doi: 10.1128/JB.00042-13.

35. De la Fuente R, Schleifer K.H., Götz F, Köst H.-P. 1986. Accumulation of porphyrins and pyrrole pigments by *Staphylococcus aureus* ssp. *anaerobius* and its aerobic mutant. FEMS Microbiol Lett 35:183–188. doi: 10.1111/j.1574-6968.1986.tb01524.x.

36. Schurig-Briccio LA, Yano T, Rubin H, Gennis RB. 2014. Characterization of the type 2 NADH:menaquinone oxidoreductases from *Staphylococcus aureus* and the bactericidal action of phenothiazines. Biochim Biophys Acta 1837:954–963. doi: 10.1016/j.bbabio.2014.03.017.

37. Gries CM, Bose JL, Nuxoll AS, Fey PD, Bayles KW. 2013. The Ktr potassium transport system in *Staphylococcus aureus* and its role in cell physiology, antimicrobial resistance and pathogenesis. Mol Microbiol 89:760–773. doi: 10.1111/mmi.12312.

38. Contreras-Moreira B, Vinuesa P. 2013. GET_HOMOLOGUES, a versatile software package for scalable and robust microbial pangenome analysis. Appl Environ Microbiol 79:7696–7701. doi: 10.1128/AEM.02411-13.

39. Epstein W. 1986. Osmoregulation by potassium transport in *Escherichia coli*. FEMS Microbiol Rev 39:73–78. doi: 10.1111/j.1574-6968.1986.tb01845.x.

40. Gründling A. 2013. Potassium uptake systems in *Staphylococcus aureus*: new stories about ancient systems. MBio 4:e00784–13. doi: 10.1128/mBio.00784-13.

41. Fung DK, Lau WY, Chan WT, Yan A. 2013. Copper efflux is induced during anaerobic amino acid limitation in *Escherichia coli* to protect iron-sulfur cluster enzymes and biogenesis. J Bacteriol 195:4556–4568. doi: 10.1128/JB.00543-13.

42. Caby M, Bontemps-Gallo S, Gruau P, Delrue B, Madec E, Lacroix JM. 2018. The EnvZ-OmpR two-component signaling system is inactivated in a mutant devoid of osmoregulated periplasmic glucans in *Dickeya dadantii*. Front Microbiol 9:2459.

43. Li YC, Chang CK, Chang CF, Cheng YH, Fang PJ, Yu T, Chen SC, Li YC, Hsiao CD, Huang TH. 2014. Structural dynamics of the two-component response regulator RstA in recognition of promoter DNA element. Nucleic Acids Res 42:8777–8788. doi: 10.1093/nar/gku572.

44. Freeman ZN, Dorus S, Waterfield NR. 2013. The KdpD/KdpE two-component system: integrating K^+^ homeostasis and virulence. PLoS Pathog 9:e1003201. doi: 10.1371/journal.ppat.1003201.

45. Fuchs S, Pané-Farré J, Kohler C, Hecker M, Engelmann S. 2007. Anaerobic gene expression in *Staphylococcus aureus*. J Bacteriol 189:4275–4289. doi: 10.1128/JB.00081-07.

46. Korotetskiy IS, Jumagaziyeva AB, Reva ON, Kuznetsova TV, Shvidko SV, Iskakbayeva ZA, Myrzabayeva AH, Ilin AI. 2019. Isolation and characterization isolates of nosocomial infections. Bul Nat Acad Sci Republic of Kazakhstan 5:199–209. doi: 10.32014/2019.2518-1467.140.

47. Korotetskiy IS, Jumagaziyeva AB, Shilov SV, Kuznetsova TV, Iskakbayeva ZA, Myrzabayeva AH, Korotetskaya NV, Ilin AI, Reva ON. 2020. Phenotypic and genotypic characterisation of clinical isolates of nosocomial infections. Eurasian J Appl Biotechnol 2020:48–60.

48. Baranyi J, Roberts TA. 1994. A dynamic approach to predicting bacterial growth in food. Int J Food Microbiol 23:277–294. doi: 10.1016/0168-1605(94)90157-0.

49. Howden BP, Beaume M, Harrison PF, Hernandez D, Schrenzel J, Seemann T, Francois P, Stinear TP. 2013. Analysis of the small RNA transcriptional response in multidrug-resistant *Staphylococcus aureus* after antimicrobial exposure. Antimicrob Agents Chemother 57:3864–3874. doi: 10.1128/AAC.00263-13.

50. Golosova O, Henderson R, Vaskin Y, Gabrielian A, Grekhov G, Nagarajan V, Oler AJ, Quiñones M, Hurt D, Fursov M, Huyen Y. 2014. Unipro UGENE NGS pipelines and components for variant calling, RNA-seq and ChIP-seq data analyses. PeerJ 2:e644. doi:10.7717/peerj.644.

51. Karp PD, Paley S, Romero P. 2002. The Pathway Tools software. Bioinformatics 18 Suppl 1:S225–232. doi:10.1093/bioinformatics/18.suppl_1.s225.

52. De Maeyer D, Renkens J, Cloots L, De Raedt L, Marchal K. 2013. PheNetic: network-based interpretation of unstructured gene lists in *E. coli*. Mol Biosyst 9:1594–1603. doi: 10.1039/c3mb25551d.

