## Supplementary material for "Comparison of transcriptional responses and metabolic alterations in three multidrug resistant model microorganisms, *Staphylococcus aureus* ATCC BAA-39, *Escherichia coli* ATCC BAA-196 and *Acinetobacter baumannii* ATCC BAA-1790, on exposure to iodine-containing nano-micelle drug FS-1": Fig. S1

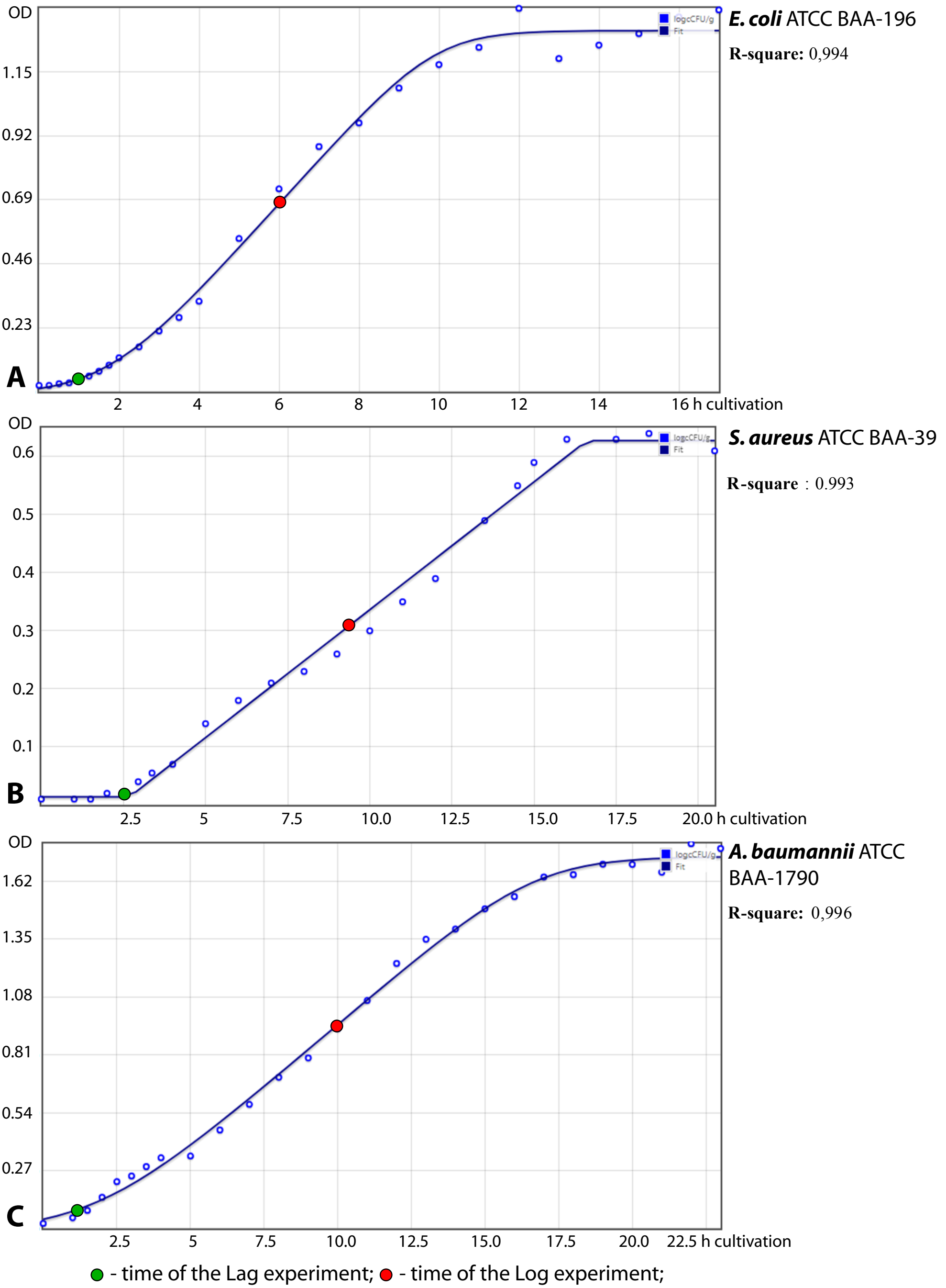


**Supplementary Fig. S1.** Growth curves of A) *E. coli* ATCC BAA-196; B) *S. aureus* ATCC BAA-39; C) *A. baumannii* ATCC BAA-1790 calculated by Baranyi and Roberts model implemented in the interactive DMFit application based on optical density (OD) values obtained in three repeats using microarray plates with MH liquid medium. Average OD values are depicted by open circles. R-square values of deviations of predicted from recorded OD values are shown.
