## Supplementary material for "Comparison of transcriptional responses and metabolic alterations in three multidrug resistant model microorganisms, *Staphylococcus aureus* ATCC BAA-39, *Escherichia coli* ATCC BAA-196 and *Acinetobacter baumannii* ATCC BAA-1790, on exposure to iodine-containing nano-micelle drug FS-1": Fig. S2

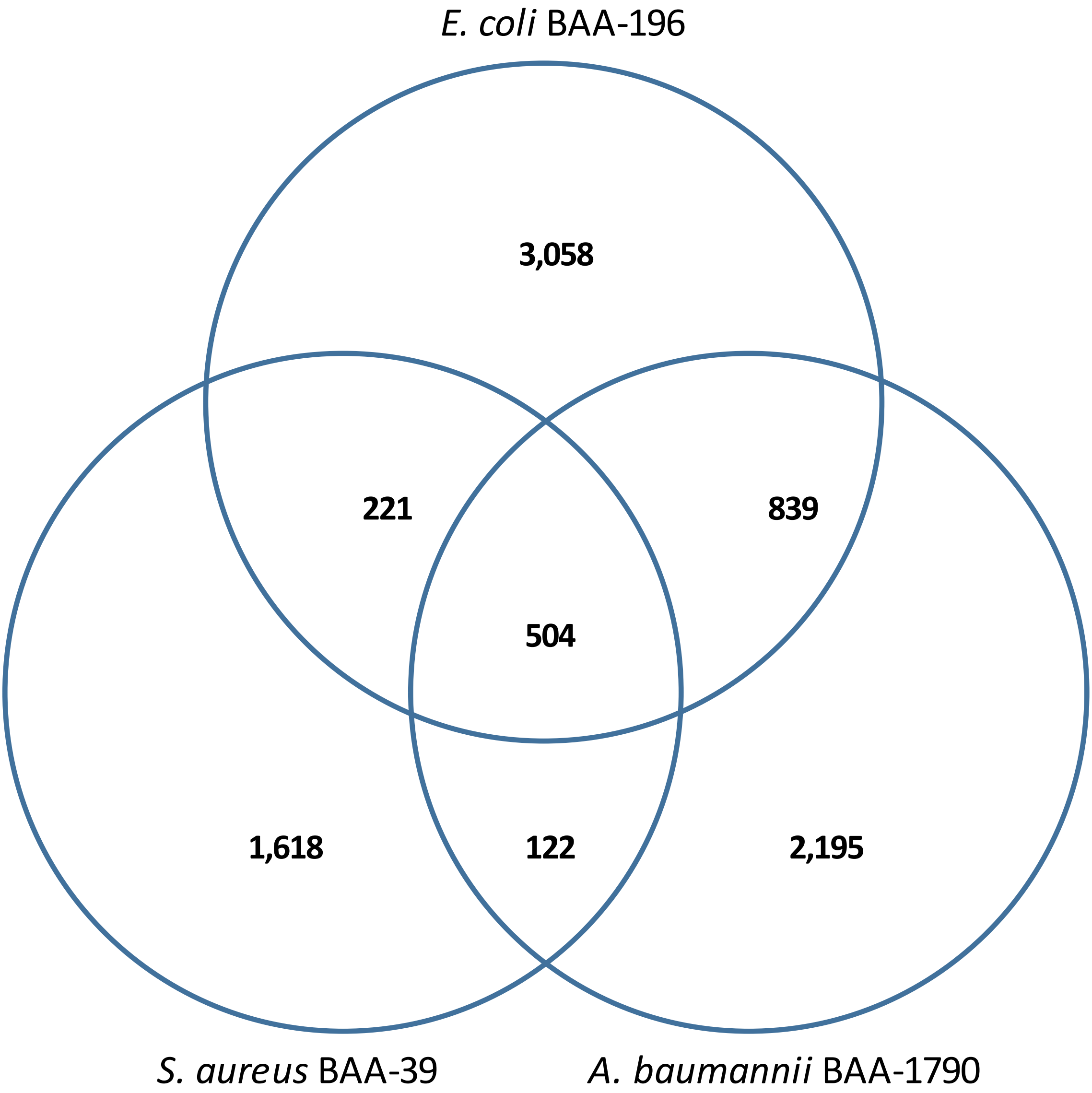


**Supplementary Fig. S2.** Venn diagram of distribution of strain-specific and homologous genes between shared by *E. coli* ATCC BAA-196, *S. aureus* ATCC BAA-39 and *A. baumannii* ATCC BAA-1790 as predicted by the program GET_HOMOLOGUES. Large admixed clusters of homologous phage-related integrases and transposases were considered as strain-specific genes.
